# Complex environmental cycles reveal evolution of circadian activity waveform and thermosensitive *timeless* splicing

**DOI:** 10.1101/2025.11.06.687109

**Authors:** Pragya Niraj Sharma, Vasu Sheeba

## Abstract

Circadian clocks synchronise physiological and behavioural rhythms to environmental cycles such as light and temperature. In nature, temperature cycles lag light cycles, with the extent of lag varying seasonally. Thus, the extent of this delay is an essential aspect of seasonal cues. Yet, the combined effects of light and temperature cycles on circadian systems remain poorly understood. Using a series of environmental cycles with varying degrees of lag between temperature and light, we examined the response of circadian activity to complex environments and show that morning and evening activity exhibit differential temperature sensitivity in *Drosophila melanogaster*. We find that the expression of temperature-induced *timeless* splice variants is modulated by light cycles as well as the degree of lag between temperature and light. We also reveal rhythmic expression of the *timeless* splicing regulator *Psi*. Furthermore, we leveraged a laboratory-selection approach to reveal that the differential temperature sensitivity of morning and evening activity evolves upon selection. We observed selection-dependent differences in temporal expression of *timeless* splice variants thus linking circadian gene splicing to behavioural plasticity. Thus, our integrated behavioural, molecular, and evolutionary approach advances the understanding of how circadian systems integrate seasonal cues.

## Introduction

Circadian clocks align physiological and behavioural rhythms with environmental cycles. This enhances fitness by enabling anticipation of daily and seasonal changes (Jabbur et al., 2024; Sharma, 2003). These biological timekeepers integrate multiple cues, like light and temperature, which vary predictably across seasons and latitudes. The duration of light (photoperiod), high temperature (thermoperiod), and the lag of temperature cycles relative to light, all vary seasonally. This is seen across latitudes, albeit with milder seasonal variation in the tropics (Ahrens & Henson, 2018). Since circadian clocks rely on both light and temperature cycles, it is crucial to understand their joint effects on circadian systems (Yoshii et al., 2010). However, most laboratory-based studies have focused on either one of these time cues in isolation.

*Drosophila melanogaster* has been instrumental in elucidating principles of circadian functioning (Dubowy & Sehgal, 2017). Most of the research has focused on light as a time cue (Helfrich-Förster, 2020) while only a few studies have focused on synchronization of circadian rhythms by temperature cycles, combined light and temperature cycles or natural conditions (Busza et al., 2007; Menegazzi et al., 2013; Yoshii et al., 2009). Existing information regarding the role of fruit fly circadian neuronal and molecular components in regulating activity patterns in response to different environmental cues makes it a useful model for studying the integration of seasonal cues (Allada & Chung, 2010; Hermann-Luibl & Helfrich-Förster, 2015). Its locomotor activity pattern is known to be modulated by both light and temperature (Dubruille & Emery, 2008) and comprises distinct morning and evening activity bouts. Each of these bouts are regulated by morning and evening neuronal subsets (M and E cells) within the Drosophila circadian pacemaker network (Grima et al., 2004; Stoleru et al., 2004). It has been demonstrated that evening activity and E cells track the temperature cycle with high fidelity when temperature rise is phase-lagged with respect to light cycle (Miyasako et al., 2007; Yoshii et al., 2010). This is not the case for morning activity or M cells which remain phase locked to lights-ON. The hypothesis that evening activity exhibits greater temperature sensitivity or temperature tracking remains to be rigorously tested since most prior studies have utilized only one phase relationship between light and temperature cycles. This results in a limited understanding of how dynamic prioritization of time cues, ‘zeitgebers’, by the circadian system can occur (Oda & Friesen, 2011). In contrast, systematic analysis of multiple phase relationships can unravel differential sensitivity of the morning and evening activity and their corresponding circadian neurons. This may underlie the encoding of seasonal cues by the circadian network, enabling optimally timed activity.

In this study, we utilized a range of phase delays of temperature cycles relative to light and systematically assayed activity patterns of flies to gain a nuanced understanding of how the circadian rhythm responds to complex zeitgeber cycles (time cues influencing the state of the circadian system). We also conducted a similar examination of activity patterns of laboratory-selected *Drosophila melanogaster* “chronotype” populations (artificially selected for divergent timing of fly emergence from pupal case for over 350 generations). This was aimed at assessing how the response of activity patterns to complex zeitgeber cycles evolves upon selection for timing of a circadian behaviour. Moreover, the overlap between the model system’s overt rhythm and underlying neuronal substrates allowed us to make inferences and develop hypotheses regarding the coupling and differential zeitgeber sensitivity between the M and E cells of the circadian neuronal network.

The coupling and connectivity across the circadian network is one of the means by which seasonal information like photoperiod or temperature is encoded into the circadian clock (Li et al., 2024; Schlichting et al., 2016, 2019; Stoleru et al., 2007). However, alternative splicing of the components of the transcription-translation feedback loop (TTFL) forms an intracellular mechanism for encoding seasonal cues. Alternative splicing of circadian genes is widespread across taxa and plays a role in maintaining robust circadian rhythms and their synchronization to time cues (Bartok et al., 2013; Shakhmantsir & Sehgal, 2019). Hence, understanding the temporal expression pattern and the role of alternative splicing of circadian genes under varying environmental conditions in flies can inform not only how a given species synchronizes to zeitgeber cycles but also help generate hypotheses which could be tested across taxa. In *D. melanogaster*, *period*, *timeless* as well as *Clock* are known to undergo temperature-dependent alternative splicing (Cai et al., 2024; Majercak et al., 1999; Martin Anduaga et al., 2019; Shakhmantsir et al., 2018). TIMELESS plays a key role in integration of both light and temperature via its interaction with the cell autonomous photoreceptor CRYPTOCHROME and temperature-dependent mRNA splicing, respectively (Dubruille & Emery, 2008; Shakhmantsir & Sehgal, 2019). A qualitative examination of expression profiles of its splice variants under temperature cycles with constant darkness versus constant light as well as across different natural cycles suggest that light and temperature cycles may have combined effects on temperature-dependent *timeless* splicing (Montelli et al., 2015; Shakhmantsir et al., 2018). While *timeless* splicing under light cycles is well studied, effects of temperature cycle and combined light-temperature cycle remain unclear (Martin Anduaga et al., 2019; Shakhmantsir et al., 2018). To address these gaps, we investigated the expression profile of *timeless* splice variants and its splicing regulator *Psi* under more complex environmental cycles.

Splice sites of circadian genes, like *period*, are known to have natural variations which are correlated with thermal adaptation, indicating that selection can shape circadian gene splicing and temperature responsiveness (Low et al., 2008, 2012). Our laboratory-selected divergent chronotype fly populations are known to exhibit differences in temperature responsiveness of their emergence as well as activity rhythms (Abhilash et al., 2019, 2020; Nikhil et al., 2014; Vaze et al., 2012). Thus, it provides an apt system to test if selection for timing of behaviour/ temporal niche can lead to divergence in *timeless* splicing and differences in circadian plasticity.

In this study, we show that morning and evening activity bouts of flies exhibit differential temperature sensitivity. Moreover, such differences depend on the phase relationship between light and temperature cycles. We further demonstrate that this differential temperature sensitivity evolves upon selection for timing of emergence. We find that the presence of light cycle along with temperature cycle and the phase relationship between the two can significantly alter temperature-dependent *timeless* splicing. We also demonstrate for the first time that the transcript of *timeless* splicing regulator, *Psi*, is rhythmic under specific environmental conditions. Furthermore, we find that selection lines with divergent emergence timing exhibit differences in the expression profiles of *timeless* splice variants. Taken together, our studies examining circadian activity rhythm under a range of environmental cycles and linking splicing profiles of a circadian gene to behavioural plasticity, advances our understanding of how circadian systems adapt to complex environmental variability.

## Results

### Temperature sensitivity of evening activity is uncovered by out-of-sync environmental cycles in *Drosophila melanogaster*

To test the hypothesis that morning and evening activity bouts differ in temperature sensitivity and to understand zeitgeber prioritization under complex environmental cycles, we conducted experiments on four wild-type *Drosophila melanogaster* populations (*control_1-4_*). First, flies were entrained to synchronized light (light: dark, LD) and temperature cycles (thermophase: cryophase, henceforth, TC) with lights-ON coinciding with temperature rise (LDTC in-sync). Subsequently, they were exposed to out-of-sync light and temperature cycles (LDTC out-of-sync). Six such experiments, ranging from 2 hr to 12 hr out-of-sync phase relationship between LD and TC, were imposed (Figure 1A). Comparison of activity profiles between LDTC in-sync and the 6 hr out-of-sync condition (4^th^ row of the schematic in Fig 1A) shows a pronounced phase shift in evening activity but not morning activity (Figure 1B). Similar activity profiles for all the phase relationships between LD and TC are shown in Figure S1A1-6.

**Figure 1.**
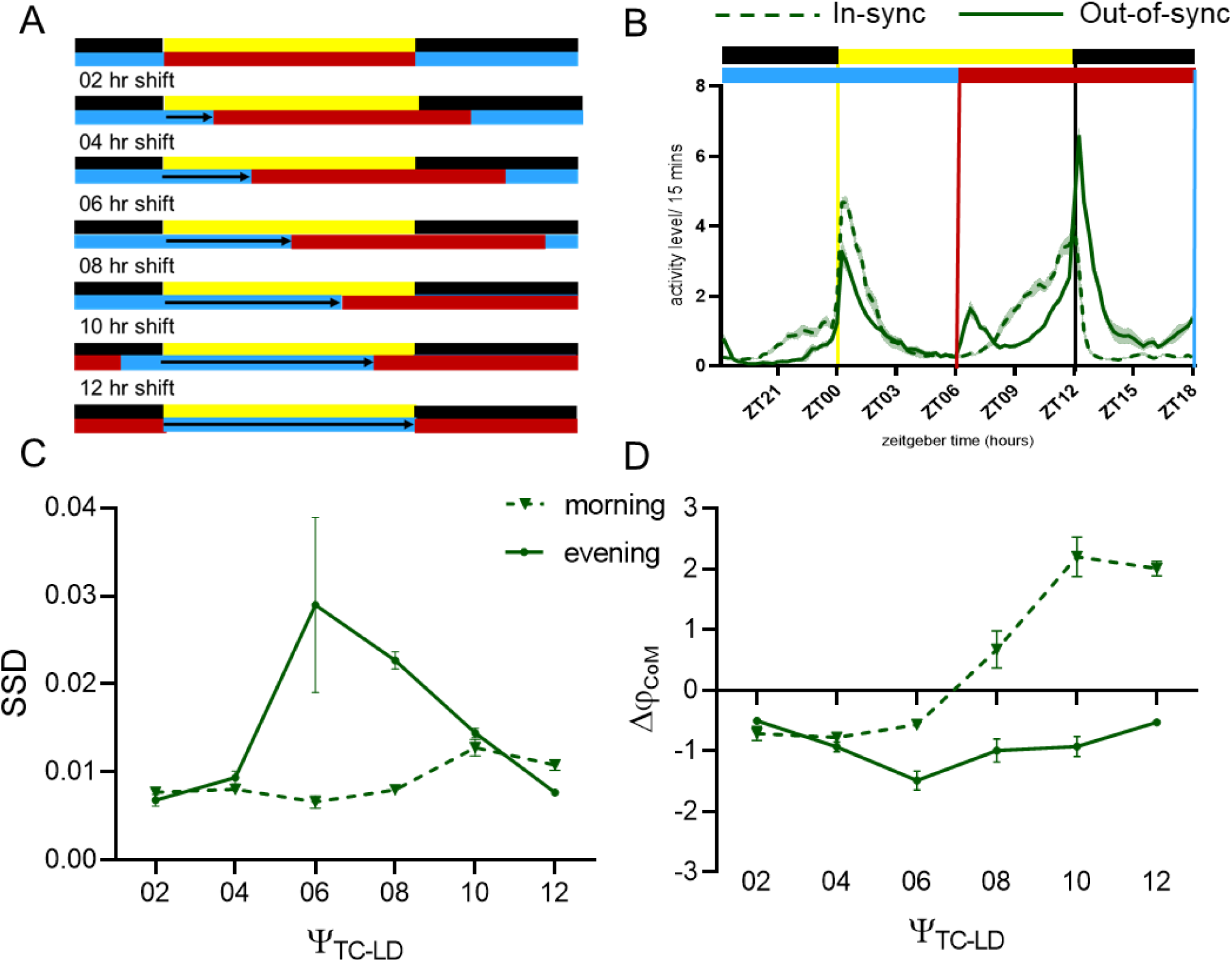
Experimental protocol and differential response of activity waveform components to out-of-sync zeitgeber cycles. (A) Schematic of all the environmental regimes. (B) Activity profiles under LDTC out-of-sync regime with TC delayed by 6 hours with respect to LD (solid trace) compared to that under LDTC in-sync regime (dashed trace). Error bands are SEM across 4 populations. For both A&B, yellow and black indicate photophase and scotophase while red and blue indicate thermophase and cryophase, respectively. Lights-ON = ZT00 (zeitgeber time 00). (C) Sum of squared differences calculated from the activity profiles under in-sync and out-of-sync zeitgeber cycles in the morning and evening window for wild-type *Drosophila melanogaster* populations for a range of phase delays of TC relative to LD (Ψ_TC-LD_). For two-way repeated measures ANOVA, time window (morning/evening) *F*_(1,18)_ = 14.75, *p* = 0.001, LDTC regime *F*_(5,18)_ = 3.588, *p* = 0.02, LDTC regime*time window *F*_(5,18)_ = 7.01, *p* = 0.0008 (D) Change in mean phase of activity (Δφ_CoM_) between in-sync and out-of-sync zeitgeber cycles in the morning and evening window for wild-type *Drosophila melanogaster* populations across a range of phase delays of TC relative to LD (Ψ_TC-LD_). For two-way repeated measures ANOVA, time window (morning/evening) *F*_(1,18)_ = 152.3, *p* < 0.0001, LDTC regime *F*_(5,18)_ = 60.52, *p* < 0.0001, LDTC regime*time window *F*_(5,18)_ = 23.72, *p* < 0.0001. Error bars are SEM across four populations of *D. melanogaster*.

The evening activity phase was delayed with the delay in TC relative to LD, whereas morning activity remained phase locked to lights-ON. We quantified these changes by comparing mean phase shifts and sum of squared differences (SSD) between synchronized and out-of-sync LDTC conditions (for details see Materials and Methods). Both metrics revealed that the degree of activity waveform change differed between morning and evening bouts and also depended on the phase difference between light and temperature (Figure 1C&D). Values for change in the mean phase of activity during the morning window are positive for 8, 10, and 12 hours of delay between the temperature and light cycles (Figure S1A4-6). It is likely that this is due to high nocturnal activity and startle activity during the morning window around the high to low temperature transition (marked by grey arrows). The negative phase shift in evening activity (Δφ_CoM_) confirmed its tracking of delayed temperature cycles (Figure 1D). Despite generally following temperature, evening activity remained aligned with light during extremely large phase delays (10–12 hours), possibly due to the highly unnatural phase relationships. These results confirm that morning and evening locomotor activities differ in temperature sensitivity, with this differential effect is contingent on the phase relationship between light and temperature zeitgebers.

Having gained a nuanced understanding of the circadian behavioural responses to complex zeitgeber cycles, we next sought to characterise the effect of environmental regimes with cyclic light and temperature on circadian gene splicing, a well-known mechanism for integrating temperature information into the molecular circadian clock. To this end, we profiled the expression of total *timeless* and all its known splice variants in one of the wild-type fly populations under three environmental conditions: (1) temperature cycles only (TC 12:12) in constant darkness, (2) synchronized light and temperature cycles (LDTC in-sync, lights-ON aligned with thermophase onset), and (3) phase-shifted light and temperature cycles (LDTC out-of-sync, thermophase onset delayed by 6 hours relative to lights-ON). We also temporally profiled *Psi*, a *timeless* splicing regulator implicated in activity phasing under temperature cycles (Foley et al., 2019).

### Light: dark cycles modulate temperature-driven *timeless* splicing

We analysed the temporal expression profile of total *timeless* using primers targeting exonic regions common to all variants. *Timeless* expression was rhythmic under TC 12:12 and LDTC in-sync but not under LDTC out-of-sync (Figure 2A1-3). Comparing TC 12:12 and LDTC in-sync revealed that the presence of a light cycle along with a temperature cycle significantly altered the *timeless* expression profile (Table S1A), consolidating expression into a sharp peak at ZT15 with increased relative amplitude (Figure 2A4, S2A). Expression profiles also differed significantly between LDTC in-sync and out-of-sync regimes, underlining the role of zeitgeber phase relationship in shaping expression and rhythmicity (Figure 2A5, Table S1B).

**Figure 2.**
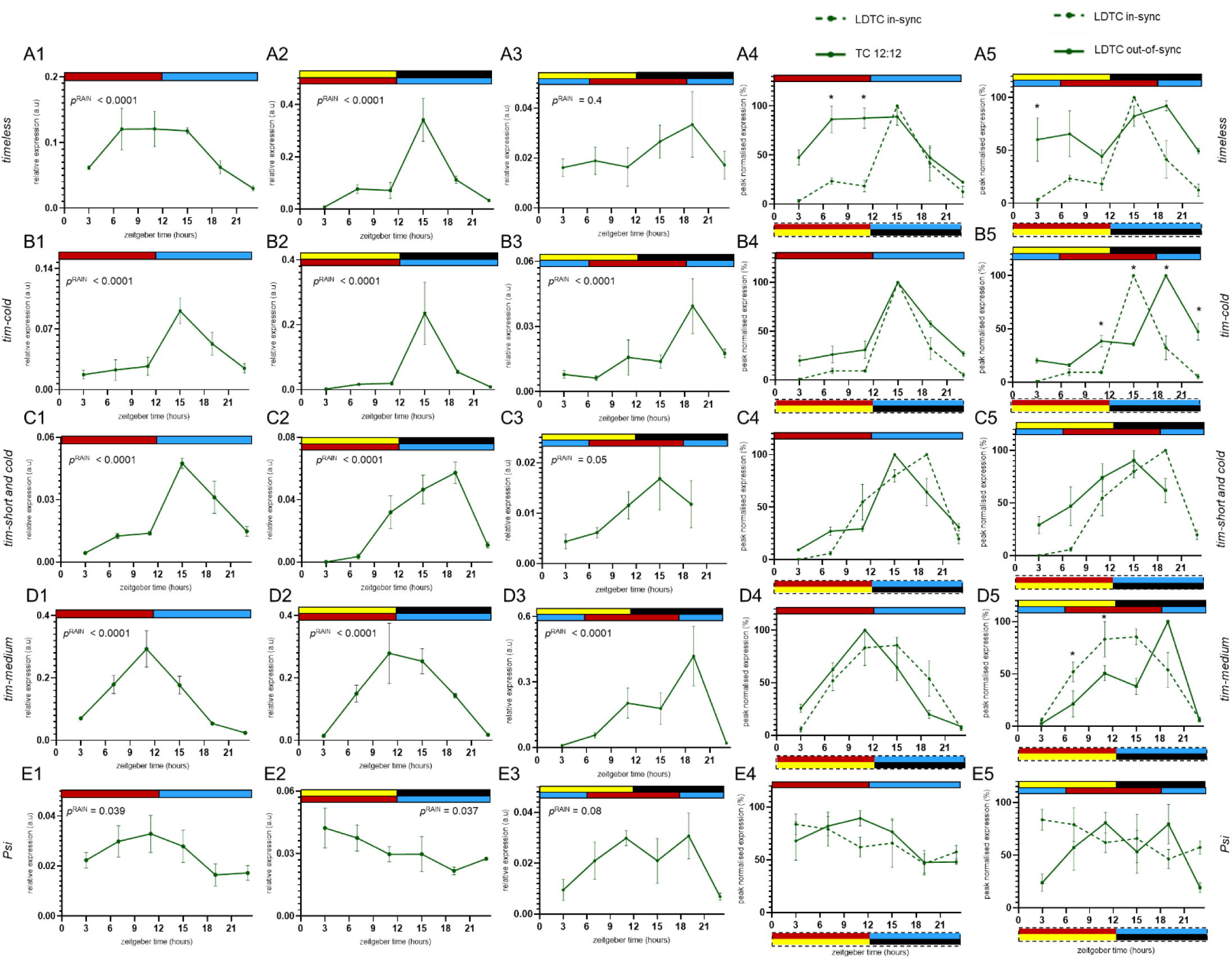
Light: dark cycle and its phase relationship with temperature cycle regulates temperature-dependent *timeless* splicing. Relative mRNA expression levels of *timeless* (A) and its splice variants (B-D) as well as splicing regulator *Psi* (E) across 24 hours sampled at 3 hr intervals. Subpanels 1-3 plot the expression levels of the indicated transcripts under the three environmental regimes – TC 12:12 (subpanels1), LD TC in-sync (subpanels 2) and LDTC out-of-sync (subpanels 3) with *p*-value for RAIN rhythmicity test mentioned inset for each. Subpanels 4 & 5 compare the peak normalised expression values between TC 12:12 versus LDTC in-sync and LDTC in-sync versus out-of-sync, respectively. Asterisks indicate significant differences at that time point between the two profiles by Sidak’s multiple comparisons test following a two-way ANOVA (with regime and time point as the factors). Colour coding for environmental cycles and ZT00 conventions are as described for Figure 1.

We next assessed temperature-induced *timeless* splice variants. The cold-induced variant *tim-cold* was rhythmic under all three regimes (Figure 2B1-3), with expression significantly altered by light presence and showing higher amplitude under LDTC in-sync compared to TC 12:12 (Figure 2B4, S2B, Table S1C). *Tim-cold*’s peak phase shifted under LDTC out-of-sync relative to in-sync, tracking cryophase onset, with significant differences in its expression profiles between these regimes (Figure 2B5, Table S1D). The other cold-induced variant, *tim-short and cold*, was rhythmic under TC 12:12 and LDTC in-sync but not under LDTC out-of-sync (Figure 2C1-3). It peaked sharply at ZT15 under TC 12:12 but under LDTC in-sync, it peaked at ZT19 with a gradual rise from ZT11. Thus, upon comparing its expression profiles under TC and LDTC in-sync, a significant regime * time point interaction and a difference in amplitude were detected (Figure 2C4, S2C, Table S1E). Its pattern was differed between LDTC in-sync and out-of-sync, suggesting that its expression profile and rhythmicity depended on light-temperature phase relationships (Figure 2C5, Table S1F). The warm temperature-induced variant, *tim-medium*, was rhythmic across all three regimes (Figure 2D1-3). Comparison between TC 12:12 and LDTC in-sync revealed that *tim-medium* expression was significantly altered upon the presence of an LD cycle along with TC (Figure 2D4, Table S1G). It showed a more gradual rise and fall under LDTC in-sync compared to the steep rise-and-fall pattern under TC 12:12 (Figure 2D4). Between LDTC in-sync and out-of-sync, the phase relationship significantly modulated its expression profile, with peak levels occurring at ZT19 under out-of-sync conditions (Figure 2D5, Table S1H).

### Rhythmic expression of *timeless* splicing regulator, *Psi*

PSI, a splicing regulator involved in *timeless* splicing, displayed rhythmic mRNA expression under TC 12:12 and LDTC in-sync but lacked statistical rhythmicity under LDTC out-of-sync (Figure 2E1-3). To our knowledge, this is the first demonstration that *Psi* exhibits environmental regime dependent rhythmicity, further linking *timeless* splicing regulation to environmental cues. While visual differences in its expression profiles across the three environmental regimes may be apparent (Figure 2E4&5), we did not detect any statistically significant differences (Table S1I&J).

Altogether our observations thus far demonstrate that the circadian system in *Drosophila melanogaster* exhibits a differential temperature sensitivity between morning and evening locomotor activity, with evening activity more closely tracking temperature cycle shifts while morning activity remains predominantly light-locked. At the molecular level, we find that the expression and rhythmicity of temperature-induced *timeless* splice variants and their regulator *Psi* are fundamentally shaped not only by ambient temperature, as previously shown, but also by light and temperature cycles as well as by the phase relationship between these environmental cues.

To examine how evolution of different circadian phases may influence the integration of light and temperature cues, we extended our behavioural and molecular analyses to laboratory-selected Drosophila populations that have been divergent phasing of their emergence rhythms. Given that it has been previously established that the circadian clocks of these populations have indeed been diverged (Abhilash et al., 2020; Kumar et al., 2007; Nikhil et al., 2016), assaying their locomotor activity patterns under a range of environmental conditions along with *timeless* splicing dynamics in these selection lines allowed us to examine how adaptive changes in the circadian system impact temperature sensitivity of the evening locomotor activity and the molecular mechanisms underlying environmental signal integration. This comparison provides critical insights into the evolutionary plasticity of circadian behaviour and its molecular regulation under complex environmental cycles.

### Evolution of temperature sensitivity: Late-selected flies exhibit enhanced evening activity phase shifts

Late-selected flies show the greatest delay in evening activity, followed by *control* flies, with early-selected flies showing the least delay (Figure 3A), while morning activity remains consistent across groups. This is especially clear in the phasing of activity drop after lights-OFF under the out-of-sync condition (indicated by grey arrows in Figure 3A). This trend among the selection lines can be observed across different phase lags between TC and LD (Figure S1A-C). We quantified timing (mean phase change) and activity pattern (SSD) changes during morning and evening periods between synchronized and each of the out-of-sync cycles. Morning activity responses were similar across the selection lines (Figures 3B, 3C), but evening activity differed distinctly (Figures 3D, 3E).

**Figure 3.**
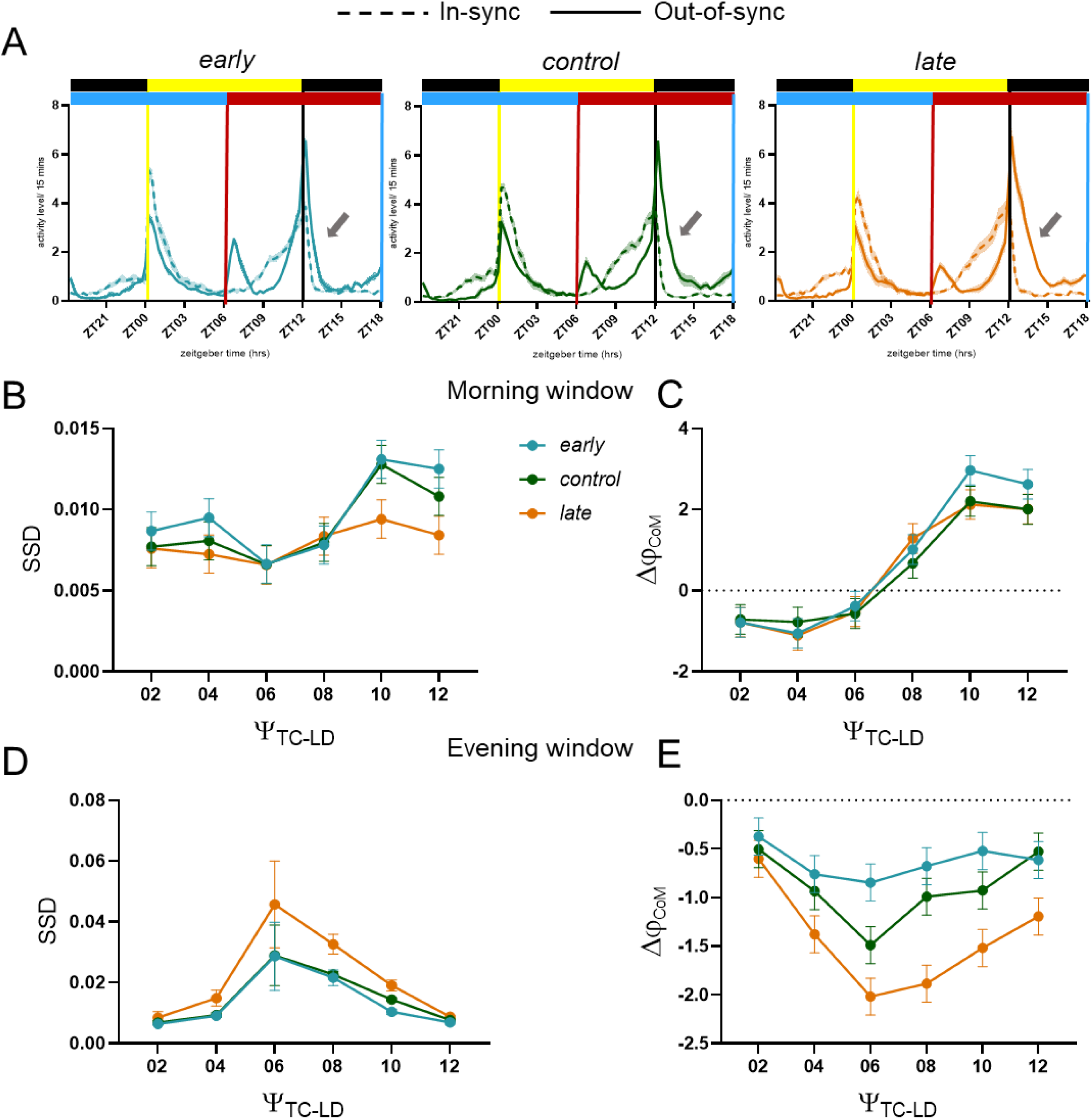
Differential sensitivity of evening activity bout among fly chronotype populations under environmental regimes with a range of phase relationship between the light and temperature cycles. (A) Activity profiles of *early_1-4_*, *control_1-4_* and *late_1-4_* selection lines under an environmental regime with light and temperature cycles in-sync (dashed trace) and out-of-sync (solid trace). Error bands = SEM across four populations. (B) Sum of square differences calculated from the activity profiles under in-sync and out-of-sync zeitgeber cycles in the morning window for the selection lines. (C) Change in mean phase of activity between in-sync and out-of-sync zeitgeber cycles in the morning window for the selection lines. (D) Sum of square differences calculated from the activity profiles under in-sync and out-of-sync zeitgeber cycles in the evening window for the selection lines with SEM error bars. (E) Change in mean phase of activity between in-sync and out-of-sync zeitgeber cycles in the evening window for the selection lines. All error bars, unless mentioned otherwise, are 95% CIs from Tukey’s HSD for the interaction effect between the fixed factors of selection and environmental regime in a three-factor ANOVA design with block as random factor, with non-overlapping error bars indicating significant differences. Colour coding for environmental cycles and ZT00 conventions are as described for Figure 1.

Mixed model ANOVA indicated significant effects of selection and TC-LD phase relationship on the change in evening activity between LDTC in-sync and out-of-sync environmental cycles (Table S2C&D), with a significant interaction indicating differential responsiveness among lines (Table S2D). Morning activity also showed interaction effects, likely due to differences in startle responses, especially when temperature changed before the morning activity bout (marked by grey arrow in Figure S1A4-6, Table S2A&B). It is clear from the mean phase change that evening activity of *late* flies tracks delayed temperature cycles to a greater extent than that of *early* or *control* flies (Figure 3E).

These findings highlight that artificial selection for timing of emergence results in distinct behavioural adjustments to environmental cues, particularly in the evening activity phase. This underscores the value of these selection lines as powerful models to investigate the correlated evolution underlying flexibility and adaptation of circadian behaviour to changing environmental conditions upon temporal niche segregation. A previous study described global differences in activity rhythms among the selection lines under TC 12:12 (Abhilash et al., 2020). Our results along with such previously established differences in temperature sensitivity of activity rhythms, make these selection lines an apt system to study correlated evolution of temperature-induced circadian gene splicing.

### Impact of long-term selection on circadian regulation of *timeless* splicing

We assayed total *timeless*, its splice variants, and the splicing regulator *Psi* in one block, i.e., *early_4_*, *late_4_*, and *control_4_* under three regimes: TC 12:12, LDTC in-sync, and LDTC out-of-sync to test whether selection drove correlated evolution in *timeless* splicing. Two comparisons were made: (1) differences among lines within each regime per transcript, and (2) changes in their expression profiles across regimes.

Total *timeless* was rhythmic under TC 12:12 in all lines, with *early_4_* showing significantly higher expression at ZT07 and ZT11 and a broader, flatter pattern in *control_4_* and *late_4_* lines (Figure 4A1). This resulted in higher relative amplitude and advanced mean phase in *early_4_* (Figure 4A4). Under LDTC in-sync, *timeless* remained rhythmic in all three lines. *Early_4_* had elevated expression at ZT11, *late_4_* had significantly lower expression at ZT15, indicating line-specific transcriptional adjustments (Figure 4A2). Under LDTC out-of-sync, rhythmicity was lost in all lines, likely due to environmental desynchrony and inherently high biological variability in our populations (Figure 4A3).

**Figure 4.**
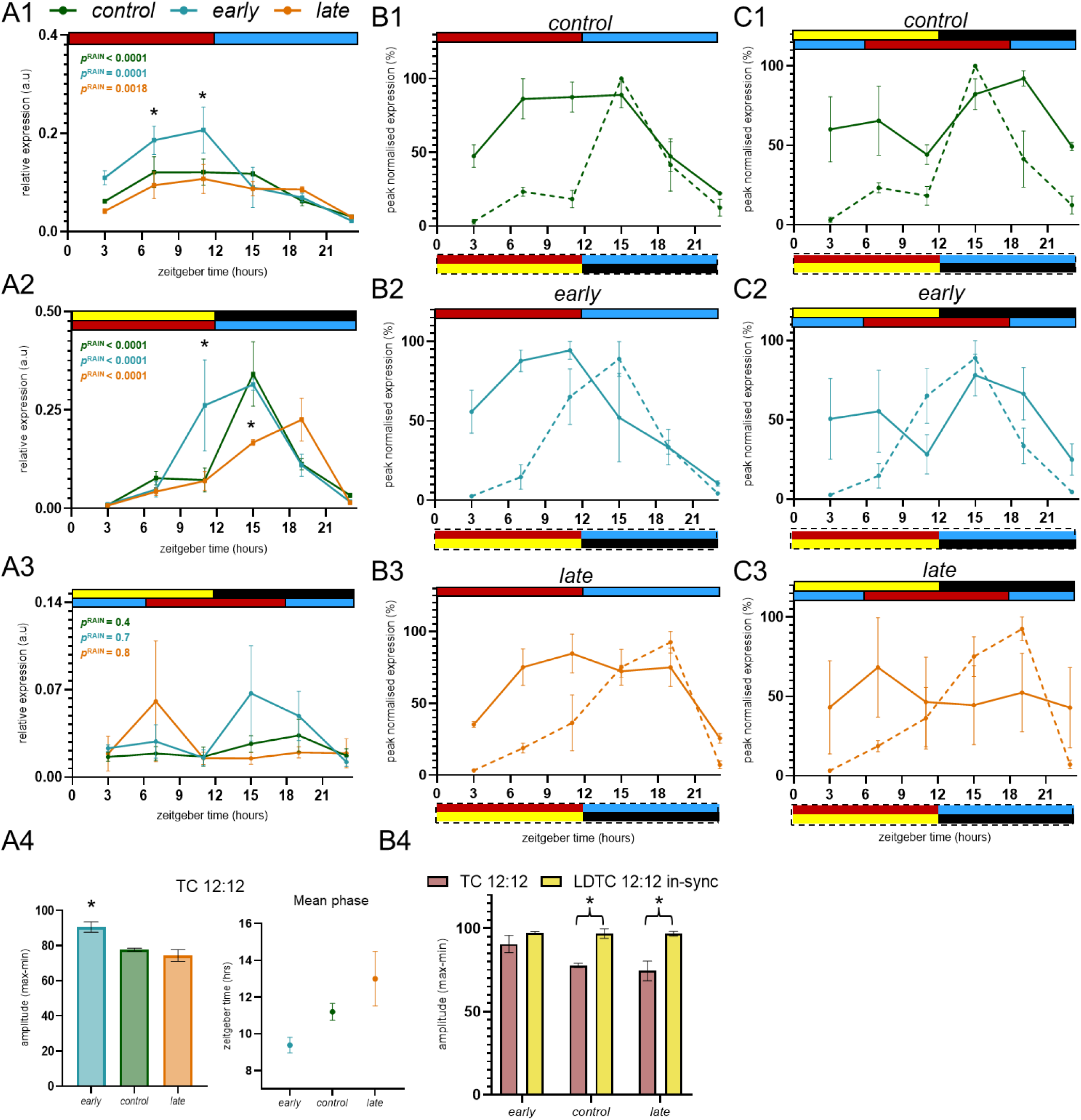
Temporal expression of *timeless* in laboratory-selected fly chronotypes across three environmental regimes. (A1-A3) *timeless* expression in one block of the selection lines *early_4_*, *control_4_*, *late_4_* under TC 12:12 (A1), LDTC in-sync (A2), and LDTC out-of-sync (A3). Asterisks indicate significant differences at those time points among the selection lines (Tukey’s HSD, two-way ANOVA). RAIN rhythmicity *p*-values inset. (A4) Amplitude (left) and mean phase (right) under TC 12:12, one-way ANOVA with Tukey’s HSD indicated by asterisk. (B1-B3) Peak-normalized *timeless* profiles in *control_4_*, *early_4_*, and *late_4_* under TC 12:12 (solid) and LDTC in-sync (dashed). (B4) Relative amplitude across environmental regimes in selection lines; one-way ANOVA with Tukey’s HSD indicated by asterisk. (C1-C3) Peak-normalized profiles under LDTC out-of-sync (solid) compared to LDTC in-sync (dashed). Environmental regimes for the solid and dashed traces are indicated at the top and bottom of the expression profiles, respectively. Colour coding for environmental cycles and ZT00 conventions are as described for Figure 1.

Comparing TC 12:12 and LDTC in-sync showed that combined light and temperature cycles delay and consolidate *timeless* expression (Figures B1-3). *Early_4_* maintained high amplitude across both, while *control_4_* and *late_4_* showed amplitude expansion under LDTC, reflecting clock plasticity (Figure 4B4). Comparing LDTC in-sync and out-of-sync revealed amplitude reduction and rhythmicity loss across lines, particularly drastic in *late_4_* flies (Figure 4C1-3). All statistical comparisons are listed in Table S3.

*Tim-cold* showed robust rhythmicity across all regimes in *early_4_*, *control_4_*, and *late_4_* (Figure 5A1-3). Under TC 12:12, *early_4_* exhibited reduced amplitude, while *late_4_* maintained higher *tim-cold* levels for longer during the cryophase. These differences in their profiles are reflected in mean phase trends and significantly higher expression in *late_4_* flies during cryophase (Figure 5A4). This correlates with previously established differences in activity phasing and temperature sensitivity among the selection lines (Abhilash et al., 2020). Under LDTC out-of-sync, peak expression was uniformly delayed to ZT19 across lines, matching delayed cryophase onset (Figure 5B1-3). Notably, at ZT11, *tim-cold* levels increased in *control_4_* and especially *late_4_* under out-of-sync compared to in-sync LDTC conditions, while *early* remained stable, revealing time-of-day dependent differential regulatory responses to environmental desynchrony (Figure 5B4).

**Figure 5.**
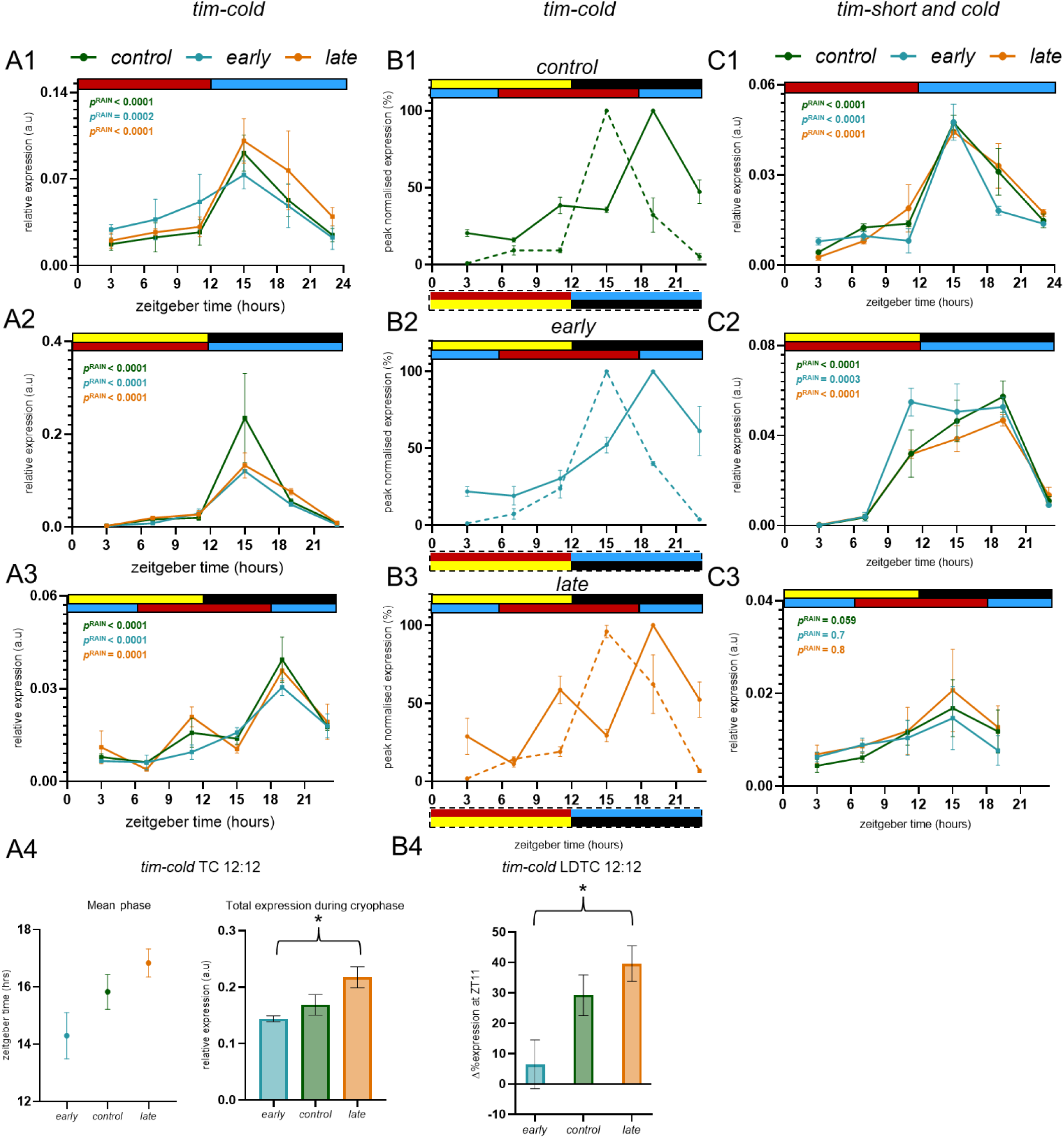
Temporal expression of cold-induced *timeless* splice variants in fly chronotypes across three environmental regimes. (A1-3) *tim-cold* expression over 24 h in selection lines under TC 12:12, LDTC in-sync, and LDTC out-of-sync, respectively; RAIN *p*-values inset. (A4) Mean phase under TC 12:12 (left) and total expression during cryophase (right) under TC 12:12; one-way ANOVA with Tukey’s HSD indicated by asterisk. (B1-3) Peak-normalized *tim-cold* expression in *early_4_*, *control_4_*, and *late_4_* lines under LDTC in-sync (dashed) and out-of-sync (solid) conditions. Environmental regimes for the solid and dashed traces are indicated at the top and bottom of the expression profiles, respectively. (B4) Change in expression level at ZT11 between out-of-sync and in-sync conditions; one-way ANOVA with Tukey’s HSD (\**p*=0.03). (C1-3) *tim-short and cold* expression across 24 h in selection lines under TC 12:12, LDTC in-sync, and LDTC out-of-sync, respectively, RAIN *p*-values inset. Colour coding for environmental cycles and ZT00 conventions as described in Figure 1. Error bars: SEM of three biological replicates.

The cold-induced *tim-sc* variant was rhythmic under TC 12:12 and LDTC in-sync but not under LDTC out-of-sync for all three lines (Figure 5C1-3). Under TC 12:12, *early_4_* showed a sharp post-peak decline at ZT19, reflecting selection line-dependent transcript modulation (Figure 5C1). Loss of *tim-sc* rhythmicity may also be contributed by increased biological replicate variability and a missing time point. All statistical comparisons for cold-induced *timeless* transcripts are in Table S4, with additional profile comparisons in Figure S3.

*Tim-medium* was rhythmic across all three lines and regimes (Figure 6A1-3). Upon comparing its expression profile under LDTC in-sync to TC 12:12 for the selection lines (Figure 6B1-3), we found that *control_4_* and *late_4_* shift in a similar manner with there being an overall delay in their profiles but significant change (Table S5D). However, *early_4_* cross 50% of their peak expression levels by ZT03 and ZT07 under TC but not under LDTC in-sync regime while the peak and decline in *early_4_ tim-medium* levels occur at the same time between the two regimes (Figure 6B2), resulting in its profile being altered significantly between these two regimes (Table S5D).

**Figure 6.**
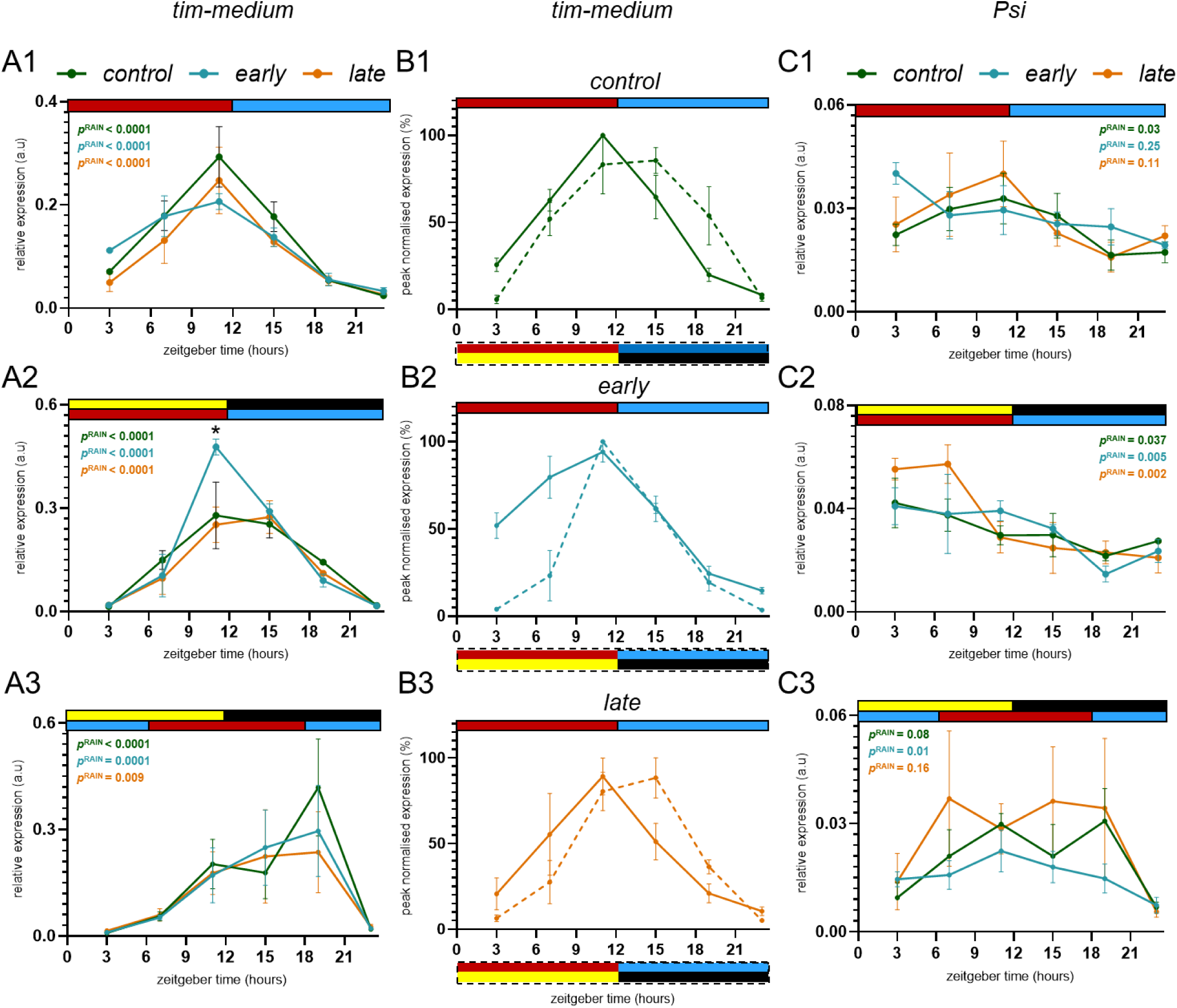
Temporal expression of warm-induced splice variant *tim-medium* and of *Psi*, a splicing regulator of *timeless*, in fly chronotype populations across three environmental regimes. (A1-3) Expression levels of *tim-medium* across a day in the selection lines under TC 12:12, LDTC in-sync regime and under LDTC 12:12 out-of-sync regime, respectively. RAIN *p*-values inset. (B1-3) Peak normalised expression profile of *tim-medium* in *control_4_*, *early_4_* and *late_4_* under TC and LDTC 12:12 in-sync conditions, solid and dashed traces, respectively. Environmental regimes for the solid and dashed traces are indicated at the top and bottom of the expression profiles, respectively. (C1-3) Expression levels of *Psi* across a day in the selection lines under TC 12:12, LDTC in-sync regime and under LDTC 12:12 out-of-sync regime, respectively. Colour coding for environmental cycles and ZT00 conventions as described in Figure 1. Error bars are SEM across three biological replicates.

*Psi* was rhythmic in *control_4_* under TC 12:12, as described before, but not in *early_4_* or *late_4_* (Figure 6C1). It was, however, rhythmic for *early_4_*, *control_4_* as well as *late_4_* flies under LDTC in-sync regimes (Figure 6C2) thus, demonstrating distinct splicing regulator dynamics linked to selection between TC versus LDTC in-sync. *Psi* was rhythmic only in *early_4_* under LDTC out-of-sync regime and not in *control* or *late* (Figure 6C3), where higher biological replicate variation may have contributed to the loss of detectable rhythmicity in addition to environmental asynchrony. All statistical comparisons conducted for *tim-medium* and *Psi* transcripts have been listed in Table S5 and all plots for profile comparisons not discussed here are a part of Figure S4.

Overall, these results detail rhythmic expression patterns and selection-dependent modulation of *timeless* splice variants and the splicing factor *Psi* across different environmental cycles. The observed variation among biological replicates and occasional arrhythmicity under LDTC out-of-sync regime in our populations could be stemming from their large standing genetic variation but appears to be compounded by the transcript being assayed as well as environmental asynchrony. This underscores both the biological complexity and sensitivity of circadian molecular regulation to environmental and genetic factors, highlighting important considerations for future studies of clock plasticity and adaptation. Altogether, our results provide strong evidence for correlated evolution of splicing dynamics of a circadian gene upon selection for timing of emergence rhythm.

## Discussion

Using an experimental protocol comprising incremental phase delays of temperature cycles relative to light, we reveal that evening activity exhibits greater temperature sensitivity compared to morning activity. Through our long-term laboratory selection approach, we find that this property of evening activity can undergo correlated evolution upon selection for divergent timing of emergence. We also demonstrate that the expression pattern of temperature-induced *timeless* splice variants is modulated by light:dark cycles. Furthermore, their expression profiles diverge upon selection for timing of emergence. We also find correlations between altered *timeless* splicing and diverged temperature sensitivity of activity rhythms among the selection lines.

### Differential temperature sensitivity of morning and evening activity

The widespread distribution of crepuscular activity across diverse lineages suggests it is an evolutionary adaptation balancing predation avoidance, light availability, and temperature regulation (Bennie et al., 2014; Carroll & Harvey-Carroll, 2023; Kirmse, 2024; Kirse et al., 2025; Rojas et al., 1999; Wong & Didham, 2024). The unparalleled advantages of the fly model system make it valuable to study how complex environmental cycles regulate crepuscular activity at molecular, neuronal and behavioural levels.

Previous studies demonstrated that in *D. melanogaster*, the molecular clock phase in evening neurons and CRY-negative neurons tracks the temperature cycle (Miyasako et al., 2007; Yoshii et al., 2009, 2010). However, such studies often used a single-phase displacement which can limit insights into the full range of circadian responses (Oda & Friesen, 2011). While Harper et al., 2016, assayed activity under a range of different phase relationships between light and temperature cycles, the temperature rise was advanced with respect to lights-ON leading to startle responses and complicating interpretation. In nature temperature rise always lags dawn, making it crucial to study the response of circadian system to phase lagged temperature cycles compared to lights-ON. Moreover, the study by Harper et al., 2016 was aimed at assessing the role of *cryptochrome* in the phasing of evening activity. Thus, the hypothesis of the two activity bouts having differential temperature sensitivity was still under-examined.

Our experimental design, incorporating incremental phase delays of the temperature cycle relative to light, enabled rigorous testing of this hypothesis under naturally plausible phase relationships between the two zeitgebers. We found that morning and evening activity bouts exhibit differential temperature sensitivity with evening activity phase tracking the delay in temperature cycle to a greater extent (Figure 1 This difference between the activity bouts was greatest under a 6-hour phase delay between the two zeitgebers, with evening activity re-aligning with the light cycle under extreme or unnatural phase lags of 10 or 12 hours (Figure 1C&D). This phase dependence underscores the importance of using incremental phase differences instead of single-phase displacement in circadian studies to avoid misleading or incomplete interpretations regarding zeitgeber integration. These behavioural dynamics under a range of phase delays between light and temperature prompt further exploration of how specific neuronal oscillators within the fly circadian network and their interactions may be mediating this phase-dependent response.

In *Drosophila melanogaster*, morning activity is regulated by PDF^+ve^ LNvs, while PDF^-ve^ small LNv and LNds govern evening activity (Allada & Chung, 2010). Integrating this cellular framework with our data and prior modelling of emergence rhythms under similar LDTC conditions (Oda & Friesen, 2011), we propose a model describing coupling and temperature sensitivity of these oscillators.

The evening oscillator group is heterogeneous (Shafer et al., 2022), containing light-sensitive neurons expressing the cell-autonomous photoreceptor CRYOTOCHROME (CRY) as well as receiving light input from morning cells and temperature-sensitive neurons lacking CRY (Beckwith & Ceriani, 2015). LNds also receive input from temperature-sensitive dorsal neurons (Li et al., 2024). Our findings support that evening and morning activity differ in temperature responsiveness, with the evening oscillator showing greater phase-dependent sensitivity. Previous work shows that fly emergence circadian rhythm too tracks the shifts in temperature albeit within a specific range before realigning to light, mirroring our observed evening activity patterns (Pittendrigh and Bruce, 1959). Another study describes the numerical simulations of limit-cycle oscillator models, which capture the main features of the emergence patterns observed under such a range of phase relationships between light and temperature cycles (Oda & Friesen, 2011). They show that such dynamics can arise from mutually coupled oscillators without asymmetry in the coupling between them or in the sensitivity to their respective dominant zeitgebers. Considering the evening oscillator’s heterogeneity and these insights, we propose a model in which distinct neuronal subpopulations integrate light and temperature inputs to regulate evening activity timing in fruit flies (Figure 7). This allows evening activity to track temperature within a defined phase range but revert to light cues under extreme phase differences, enabling seasonal adjustment through modulation of oscillator phase relationships. Beyond modelling the observed activity data, certain aspects of this model can be experimentally tested by using cell-specific knockout or knockdown of synchronizing factors such as the neuropeptide pigment dispersing factor (PDF) or of photoreceptor *cryptochrome*, and assessing their effects on activity under LD-TC conditions.

**Figure 7.**
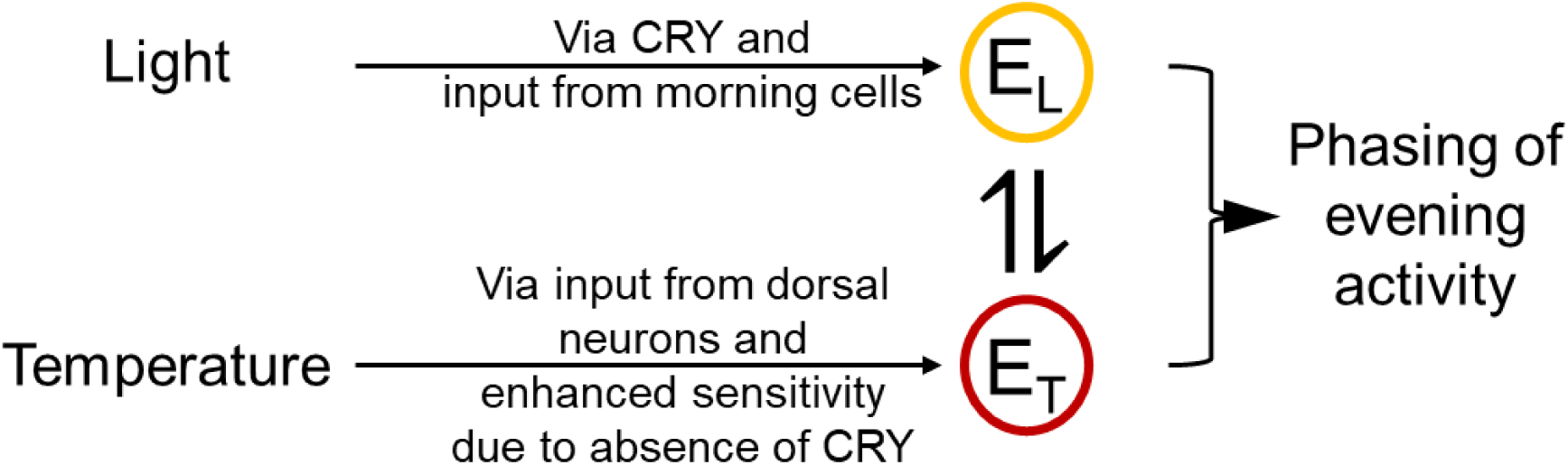
Model for integration of light and temperature signals by evening oscillator neurons to fine tune the timing of evening activity: The evening oscillator cluster consists of two mutually coupled neuronal groups—one predominantly light-sensitive (E_L_) and the other predominantly temperature-sensitive (E_T_). The phasing of evening activity results from their combined influence in response to environmental cues. Photic input affects the light-sensitive group via its coupling with morning cells and presence of CRY, while temperature input influences the temperature-sensitive group through its connection with temperature-sensitive dorsal neurons and absence of CRY. Additionally, the overall evening activity phase depends on the coupling between E_L_ and E_T_. This model assumes symmetric sensitivity to zeitgebers and mutual coupling between E_L_ and E_T_, consistent with theoretical models for emergence rhythms under similar experimental conditions.

### *timeless* splicing as a node for integration of seasonal cues

In addition to neuronal circuit mechanisms, molecular pathways such as temperature-dependent alternative splicing of the *timeless* gene offer a complementary route for integrating seasonal cues into the circadian system. It is well established that *timeless* undergoes temperature-dependent splicing: cooler temperatures upregulate *tim-cold* (*tim-c*) and *tim-short and cold* (*tim-sc*), while warmer temperatures increase *tim-medium* (*tim-M*) (Martin Anduaga et al., 2019; Shakhmantsir et al., 2018). These prior studies have primarily examined *timeless* splicing under LD cycles with constant cool or warm temperatures but without precise quantitative comparisons across environmental conditions with cyclic temperature and cyclic light.

Our study systematically profiled total *timeless* and its splice variants under three distinct environmental conditions (i) TC – temperature cycle under constant darkness, (ii) LDTC in-sync – synchronous light and temperature cycle (iii) LDTC out-of-sync – temperature rise was delayed by 6 hours with respect to lights-ON. We found that *timeless* expression was rhythmic under TC and LDTC in-sync (Figure 2A1&2). However, the amplitude and consolidation were higher under LDTC in-sync, reflecting synergistic effects of synchronous environmental cues (Figure 2A4, Figure S2A). In contrast, LDTC out-of-sync regime delayed and dampened *timeless* expression highlighting the critical effect of phase alignment between zeitgeber cycles on *timeless* rhythmicity (Figure 2A5).

The cold induced isoform, *tim-cold*, showed robust rhythmicity across all regimes (Figure 2B1-3). While *tim-cold* expression profile was significantly altered by the presence of light cycle and its phase relationship with temperature cycle (Figure 2B4&B5, Table S1C&D), its peak expression was always phase locked to cryophase onset. However, the other cold-induced transcript, *tim-sc* showed more complex dynamics, it peaked with *tim-cold* only under TC but had a gradual rise and a delayed peak under LDTC in-sync (Figure 2C1&2). This suggests that its peak expression is not universally locked to temperature drop. Its expression pattern under TC and LDTC in-sync suggest that *tim-sc* may act as a seasonal detector, integrating both light and temperature information (Figure 2C4, Table S1E). On the other hand, *tim-cold* may relay immediate temperature drop information to the circadian clock. Our hypothesis is supported by the presence of *tim-sc* in temperate *D. simulans* and *D. melanogaster* and its absence in tropical *D. yakuba*, while *tim-cold* is expressed in all three species (Martin Anduaga et al., 2019). Additionally, *tim-cold* likely binds AGO1-miRNA complexes, whereas TIM-SC protein regulates PERIOD stability, highlighting functional differences between these cold-induced isoforms (Martin Anduaga et al., 2019).

The warmth-induced transcript *tim-medium* is under strong post-transcriptional regulation by the RNAi machinery, and the protein of this transcript is not functional (Martin Anduaga et al., 2019; Shakhmantsir et al., 2018). Therefore, it is thought to contribute to delays in TIM protein profile under warmer temperatures by effectively reducing the levels of the canonical *tim-long* variant available for translation (Shakhmantsir et al., 2018). In our study, the rhythmic expression of *tim-medium* across all environmental conditions highlights its robust regulation (Figure 2D1-3). Furthermore, the significant alterations in its expression profile caused by the presence of a light cycle and the phase relationship between light and temperature underscore the complex interplay of these cues in likely modulating TIM protein levels via the expression dynamics of *tim-medium* (Figure 2D4&5, Table S1G&H).

Critical to *timeless* alternative splicing are the splicing factors PSI and PRP4, which likely regulate *timeless* splicing in opposite directions: *prp4* knockdown upregulates *tim-medium* while *Psi* knockdown downregulates it but increases the cold-induced isoforms under LD cycles with constant temperature (Foley et al., 2019; Shakhmantsir et al., 2018). Due to *Psi*’s role in activity phasing under TC we conducted time point profiling of this splicing regulator under three environmental conditions (Foley et al., 2019). We show for the first time that *Psi* expression is rhythmic under certain environmental conditions (Figure 2E1&2).

Previous work suggests a positive correlation between *Psi* and *tim-medium* levels and a negative correlation between *Psi* and levels of *tim-cold* and *tim-sc* (Foley et al., 2019). We observe that this correlation is most clearly upheld under temperature cycles wherein *tim-medium* and *Psi* peak in-phase while the cold-induced transcripts start rising when *Psi* falls (Figure 8A1&2). Under LDTC in-sync conditions this relationship is weaker, even though *Psi* is rhythmic, potentially due to light-modulated factors influencing *timeless* splicing (Figure 8B1&2). This is in line with *Psi*’s established role in regulating activity phasing under TC 12:12 but not LD 12:12 (Foley et al., 2019). This is also supported by the fact that while *tim-medium* and *tim-cold* are rhythmic under LD 12:12 (Shakhmantsir et al., 2018), *Psi* is not (datasets from Kuintzle et al., 2017; Rodriguez et al., 2013). Assuming this to be true across genetic backgrounds and sexes and combined with our similar findings for these three transcripts under LDTC out-of-sync (Figure 2B3, D3&E3), we speculate that *Psi* rhythmicity may not be essential for *tim-medium* or *tim-cold* rhythmicity especially under conditions wherein light cycles are present along with temperature cycles (Figure 8C1&2). Our study opens avenues for future experiments manipulating PSI expression under varying light-temperature regimes to dissect its precise regulatory interactions. We hypothesize that flies with *Psi* knocked down in the clock neurons should exhibit a less pronounced shift of the evening activity under the range LDTC out-of-sync conditions for which activity is assayed in our study. On the other hand, its overexpression should cause larger delays in response to delayed temperature cycles due to the increased levels of *tim-medium* translating into delay in TIM protein accumulation.

**Figure 8.**
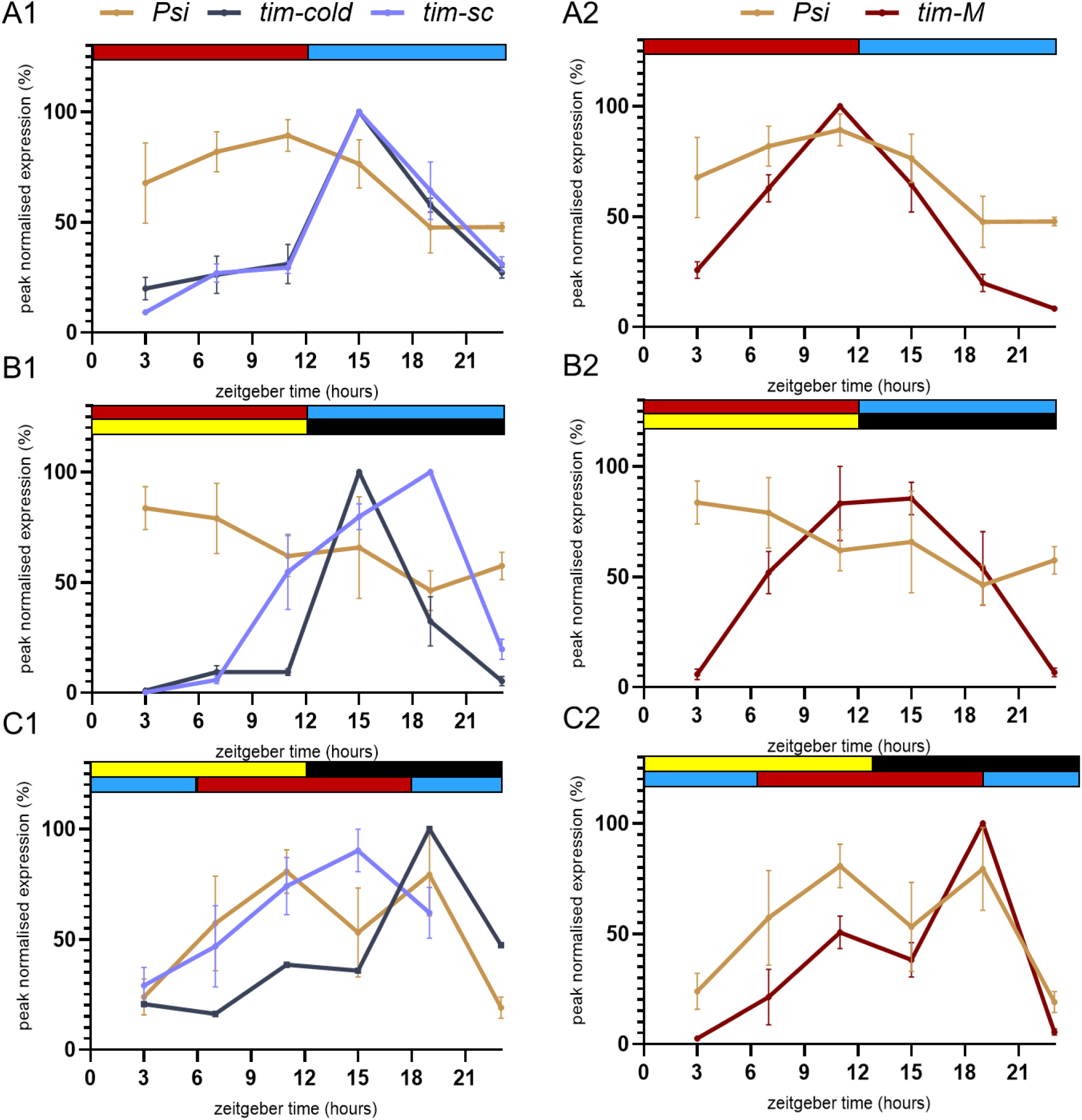
Expression pattern of *Psi* along with the warm and cold-induced transcripts in *control_4_* under three environmental regimes. (A1&2) Peak normalised expression profile of *Psi* with the cold-induced transcripts and *tim-medium* under TC 12:12. (B1&2) Peak normalised expression profile of *Psi* with the cold-induced transcripts and *tim-medium* under LDTC in-sync. (C1&2) Peak normalised expression profile of *Psi* with the cold-induced transcripts and *tim-medium* under LDTC out-of-sync. Lights-ON = ZT00 (zeitgeber time 00). Colour coding for environmental cycles and ZT00 conventions as described in Figure 1. All error bars are SEM across three biological replicates.

Our detailed characterisation and comparison of *timeless* splicing across environmental regimes places *timeless* splicing as a pivotal node in seasonal cue integration since it is influenced not only by ambient temperature but also by light cycle and its phase relationships with temperature cycle. Our study underscores a complex interplay of environmental signals in regulating activity patterns as well as circadian gene splicing.

Another key aspect of seasonal variation is photoperiod; since *period* splicing is sensitive to photoperiodic conditions, *timeless* splicing may be as well (Majercak et al., 2004). Previous studies have characterised naturally occurring variation in splice site strength in the circadian gene *period* along with clinal studies of its SNPs and its correlation with seasonal adaptation highlights the evolutionary importance of circadian gene splicing (Low et al., 2008, 2012). Such clinal studies for splice site strength within *timeless* in *D. melanogaster* could provide insights into their role in adaptation to varying environmental cycles. While clinal studies are indispensable for studying trait evolution, including that of circadian rhythms, they are largely correlational and ill-equipped to isolate the influence of specific environmental factors (Abhilash & Sharma, 2016). On the other hand, laboratory selection experiments, while lacking ecological complexity, address these limitations by focusing on a specific selection pressure. These provide advantages such as no assumptions regarding ancestry and ancestral state of the trait of interest, replicate populations, stronger causal inference regarding the effect of selection as well as controlled population genetics (Abhilash & Sharma, 2016). Therefore, through our approach of using laboratory populations selected for divergent timing of fly emergence, we could investigate how circadian system adaptions influence evening activity temperature sensitivity and *timeless* splicing and gain insights into evolution of seasonal cue integration under a temporal niche segregation-like scenario.

### Correlated evolution of temperature sensitivity of evening activity and circadian gene splicing upon selection for timing of emergence

Late-selected flies diverged in the temperature sensitivity of their evening activity, with their evening activity phase tracking the shift in temperature cycle to a greater extent than that of *early* and *control* flies (Figure 3). We hypothesise that under out-of-sync LDTC conditions, PERIOD and TIMELESS expression in CRY-negative neurons will shift most in *late* populations, mirroring evolved plasticity of evening activity in response to temperature cues. The divergence observed in *late* flies taken together with our findings for *control* (wild-type populations) indicate that differential temperature sensitivity of morning and evening activity bouts can undergo correlated evolution upon selection for timing of circadian emergence. Such behavioural dynamics in our selection lines make it an apt system to explore how our circadian organisation model proposed for *control* (Figure 7) may have been altered upon selection. We propose a model for the circadian organisation in *late* which can explain the behavioural pattern observed (Figure 9). We propose testable simplistic parsimonious models considering our own observations, known connectivity across the circadian network, and the findings from a previous study (Oda & Friesen, 2011).

**Figure 9.**
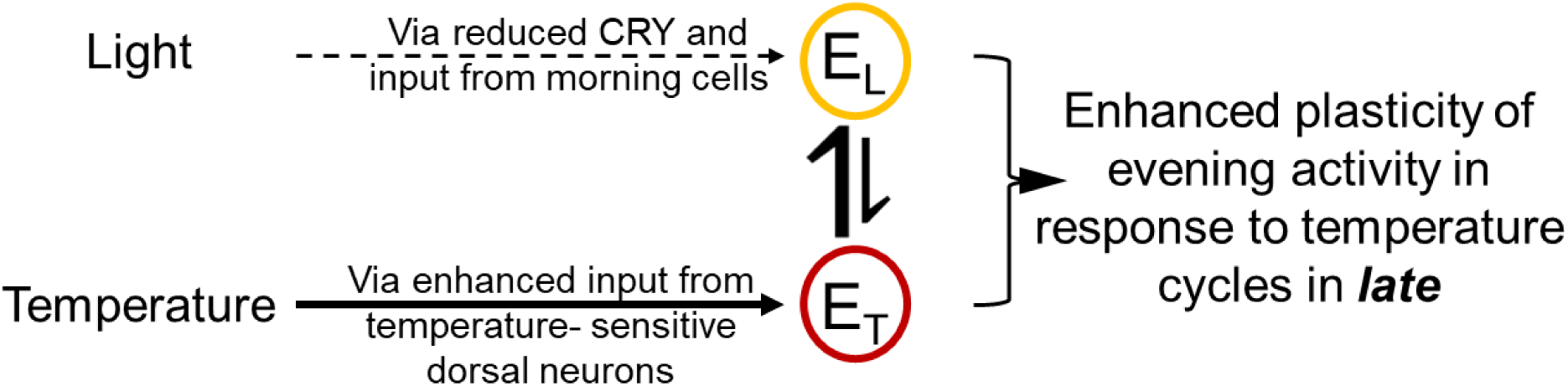
Model for correlated evolution of enhanced plasticity of evening bout in *late* populations in response to temperature cycles: Any one or more of the above mechanisms could be involved in the enhanced plasticity of evening activity in *late* in response to temperature cycles. *Late* flies show reduced levels of the *cry* transcript (Nikhil et al., 2016), lower CRY can facilitate a stronger response to temperature cues (Harper et al., 2016). This is also possible if the evening oscillator receives weaker input from the morning cells and stronger input from the temperature-sensitive dorsal cells in *late* populations compared to *control*. Finally, asymmetry in coupling between the E_L_ and E_T_ in the above indicated direction can also cause stronger responses to temperature cues because E_T_ could then dictate the rhythms in E_L_.

Building on the behavioural divergence observed, we investigated whether *timeless* splicing plays a role in this adaptation by examining molecular correlates that might underlie differences in temperature sensitivity among the selection lines. Under TC 12:12, *timeless* expression is advanced in *early_4_* and delayed in *late_4_* relative to *control* flies (Figure 4A1&A4), mirroring their activity rhythm phases (Abhilash et al., 2020). Mathematical models propose that a larger amplitude oscillator is predicted to have reduced resetting capacity, whereas smaller amplitude oscillators reset more readily (Barrett & Takahashi, 1997; Winfree, 2001). Overexpression of clock components reduces oscillator amplitude and increases resetting capacity (Busza et al., 2007), making the clock more responsive to environmental perturbations (Pittendrigh et al., 1991). *Early_4_* having the highest amplitude of *timeless* expression (Figure 4A4) aligns with them taking the longest to re-entrain activity rhythms after jetlagged temperature cycles, while *late* flies re-entrain fastest (Abhilash et al., 2020). Since *late_4_* and *control_4_* flies do not differ in *timeless* amplitude, other factors likely contribute to the rapid resetting under temperature cycle jetlag shown by *late* flies compared to *control*, perhaps reflecting previously described divergence in their circadian organisation (Figure 7 vs. Figure 9).

The most striking differences in *timeless* splicing among the selection lines were observed for *tim-cold* and the warm temperature-induced *tim-medium* transcripts, both rhythmic across all three lines and environmental conditions (Figure 5A1-3, Figure 6A1-3). *Tim-cold* transcript levels were significantly elevated in *late_4_* flies during cryophase under TC 12:12 (figure 5A4), consistent with their heightened temperature responsiveness (Abhilash et al., 2020). Comparing TC and LDTC in-sync regimes suggests that LD modulates *tim-cold* expression differentially across selection lines (Figure 5B1-3). The phase relationship between light and temperature cycles (LDTC out-of-sync vs. in-sync) also impacts *tim-cold*, with *late_4_* flies showing the largest change at ZT11 (Figure 5B4). However, linking these molecular shifts to behaviour mandates mutant or transgenic studies targeting *tim-cold*. While *tim-medium* expression profile was significantly altered when LD was present along with TC in *early_4_*, this was not the case for *control_4_* and *late_4_* flies (Table S5D, Figure 6B1-3). Thus, not only is *tim-medium* expression profile altered in response to LD in addition to TC in wildtype flies (*control_4_*) as mentioned before, but this response diverged upon selection for timing of emergence rhythm. Under LDTC out-of-sync, *tim-medium* accumulation was delayed, peaking at ZT19 in *early_4_* and *control_4_* lines (Figure S4A1&2), whereas *late_4_* flies maintained high expression from ZT11 to ZT19, broadening the profile (Figure S4A3). Advanced *tim-medium* rise is known to delay TIM protein accumulation and thus would delay activity (Shakhmantsir et al., 2018). Accordingly, *tim-medium* expression under LDTC out-of-sync aligns with the delayed evening activity observed in *late_4_* flies (Figure 3A). Taken together, these findings strongly indicate that *timeless* splicing dynamics evolve under selection for divergent timing of fly emergence. However, our molecular conclusions are currently limited to one population per selection line, unlike our behavioural data drawn from four populations for each selection line.

Our study further reveals distinct rhythmicity of *Psi*, a *timeless* splicing regulator, among *early_4_*, *control_4_*, and *late_4_* flies, highlighting selection’s impact on this splicing regulator. *Psi* rhythmicity is disrupted in *early_4_* and *late_4_* flies under temperature cycles but maintained in *control_4_* (Figure 6C1). Under synchronized LDTC, *Psi* is rhythmic across all lines (Figure 6C2). Under desynchronized LDTC, rhythmicity is largely confined to *early_4_* flies, with high variance suggesting complex gene-environment interactions shaping its expression (Figure 6C3). Given PSI’s role in neural splicing and developmental processes (Labourier et al., 2002), it is plausible that *Psi* has been a direct target of selection for emergence timing.

This raises a compelling question: What functional roles do *timeless* splice variants serve in emergence rhythms, which are regulated by the same central clock as activity rhythms (Qiu & Hardin, 1996)? Emergence rhythms of flies deficient in specific splice variants would illuminate how seasonal signals and environmental temperatures influence holometabolic development via the circadian clock. Our selection lines too, provide an ideal system to investigate this question, as they exhibit divergent temperature sensitivity in emergence rhythms as well (Abhilash et al., 2019; Nikhil et al., 2014). Since introgressing transgenic constructs into these backgrounds may be challenging, RNA sequencing of brains from freshly emerged flies—particularly under TC or LDTC out-of-sync regimes—could reveal variant-specific expression relevant to differential temperature sensitivity of emergence rhythms. Multiple studies conducted across generations on these laboratory-selected fly populations, including the current one, consistently demonstrate that selection for emergence timing using light as the sole cue leads to divergence in temperature sensitivity of both emergence and activity rhythms.(Abhilash et al., 2019, 2020; Nikhil et al., 2014; Vaze et al., 2012). These studies also highlight that laboratory selection has uncovered novel and unexpected genetic correlations, such as between emergence timing and temperature responsiveness of circadian rhythms. Through the current study we reinforce the notion that selection for behavioural phasing fundamentally reshapes the neuronal network and molecular circadian machinery underlying these rhythms.

In summary, our findings demonstrate that the temperature sensitivity of evening activity depends on the degree of misalignment between the light and temperature cycles; most likely due to the heterogeneous nature of evening oscillator neurons. This highlights how circadian systems can dynamically prioritize different environmental signals, a principle relevant to understanding how organisms cope with shifting environmental features or seasonal changes. Our detailed profiling of *timeless* splice variants under different light and temperature regimes revealed that in addition to ambient temperature, *timeless* splicing is modulated by the presence of light cycles along with temperature cycles and the phase relationship between the two. This shows that alternative splicing of core clock genes is a key mechanism for integrating multiple time cues, a process which may likely be conserved across taxa. Using the laboratory-selection lines, we reveal that selection for emergence timing leads to correlated evolution of temperature sensitivity of evening activity and *timeless* splicing. Our findings show how differential temperature response of the activity components can provide a means to fine tune behaviour under complex environmental cycles. Additionally, we highlight how *timeless* splicing is dynamically regulated by these cycles and how selection shapes both behaviour and circadian gene splicing. Altogether these findings emphasize the circadian clock’s complexity and adaptability.

## Materials and Methods

### Fly strains

#### Wildtype fly populations and laboratory-selected divergent chronotype fly populations

Four large and outbred populations were used to derive four of each *early*, *control* and *late* selection lines. Therefore, the set of selection lines comprises 12 populations – *early_1-4_*, *control_1-4_* and *late_1-4_*. The *early*, *control* and *late* selection lines with the same subscript indicate that they have been derived from the same ancestral population (referred to as ‘blocks’). In contrast, different subscripts indicate independent genetic substructures. The *early* and *late* selection lines have been subjected to selection for phase of entrainment of adult fly emergence for more than 400 generations now while the *control* populations remain unselected, comprising of flies emerging throughout the day and represent wildtype flies (detailed maintenance protocol is described in Kumar et al., 2007). It is important to note that the wildtype *control* or the selected *early* and *late* are reared in large numbers (∼1200 flies in roughly 1:1 sex ratio for each population) and therefore, have high standing genetic variation.

One generation of common rearing wherein no selection pressure is applied and flies emerging throughout the day are collected to form the parental generation from which egg collection is conducted for the experimental flies. This minimises the maternal and non-genetic inheritance effect on the measured trait (Bonduriansky & Day, 2009).

All the blocks of *early*, *control* and *late* have been used to test for the differential responses of the activity rhythm under environmental regimes with light and temperature cycles. Below is the table listing the number of generations of selection these populations had undergone at the time when each of these activity assays were performed.

| LDTC regime (LDTC - lag between TC and LD in hours) | Number of generations of selection |
| --- | --- |
| LDTC - 2 hrs | 366 |
| LDTC - 4 hrs | 363 |
| LDTC - 6 hrs | 366 |
| LDTC - 8 hrs | 363 |
| LDTC - 10 hrs | 365 |
| LDTC - 12 hrs | 365 |

#### Locomotor activity recording

3-5 days old virgin male flies from each of the 12 populations (4 blocks of each of the three selection lines post common rearing, *early_1-4_*, *control_1-4_* and *late_1-4_*) were housed in locomotor tubes with corn food medium and placed in Drosophila Activity Monitors (DAM system by Trikinetics) to measure their activity counts. Assays were carried out in Sanyo incubators (MIR-154, Sanyo, Tokyo, Japan). Light intensities were adjusted to 70 lux using 298 neutral-density Lee filter and a LI-COR light meter. The maintenance of these populations is carried out under 70 lux, and light intensities lower than 100 lux minimise startles. The temperature cycle used had a warm/thermophase temperature of 28 °C and a cool/cryophase temperature of 19 °C. For each of the six experiments in which TC lagged LD, the activity was first recorded under conditions wherein LD and TC were in-sync and were transferred to one of the LDTC out-of-sync regimes after 4 days. This allowed for calculations to compare the waveform under the LDTC out-of-sync regime to that of LDTC in-sync regime from the same generation/set of flies for each experiment.

**Figure 10.**
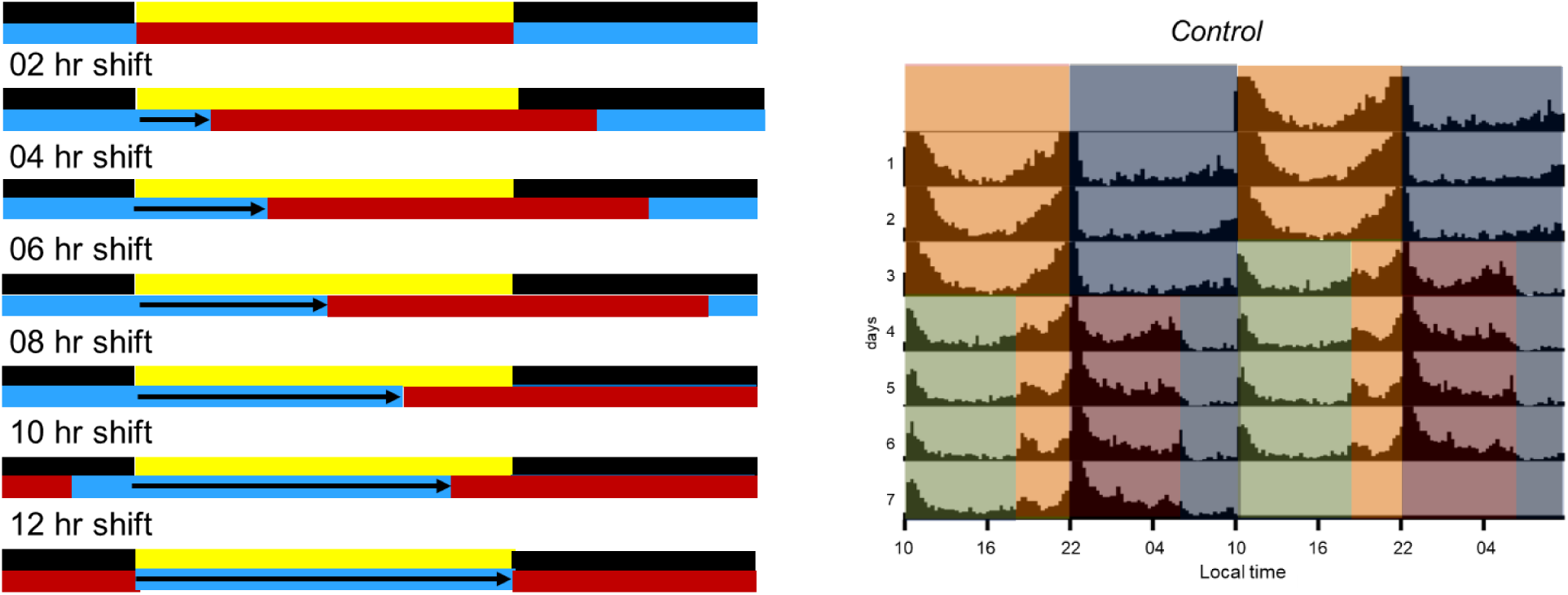
Experimental protocol. (left) schematic of all the environmental regimes used in this study and (right) an example batch actogram, activity was first recorded under conditions with light (70 lux) and temperature cycles (19 °C and 28 °C) being in-sync and then transferred to one of the out-of-sync regimes. In this example flies were transferred on the 4^th^ day of recording to an environmental regime wherein the temperature cycle was delayed by 8 hours with respect to the light cycle. Green hue = light + low temperature, orange hue = light + high temperature, red hue = darkness + high temperature and grey hue = darkness + low temperature.

#### Activity profile analyses

The activity profiles for each individual under the in-sync and out-of-sync regimes were calculated by averaging the daily profiles under each of the regimes. When a set of flies were transferred from the in-sync regime to an out-of-sync regime, the first couple of cycles were eliminated from the calculation for average profiles since all flies attained stable phases after day 2 under the new LDTC regime. The activity counts in average profile for each individual under both in-sync and out-of-sync regime were normalised by the total average activity from that individual to obtain activity level per 15 minutes. These two profiles for each individual were then used to assess changes in the activity waveform between the two regimes.

#### Assessing phase lability and changes in activity waveform under LDTC out-of-sync conditions compared to LDTC in-sync conditions

The activity profiles were divided into a 12-hour morning window (6 hours before and after lights-ON) and an evening window (6 hours before and after lights-OFF). The following calculations were conducted for the activity within each of these time windows without filtering the data or elimination of startle peaks to conduct unbiased assessment of the response of the morning and evening activity components.

##### Sum of squared differences (SSD)

The activity level in a particular 15-minute bin under the LDTC out-of-sync regime was subtracted from the activity level in the same 15-minute bin under the LDTC in-sync regime of the same individual. This difference within a particular bin was then squared. These squared values were then summed up over the 12 hour window to give an SSD value for an individual. SSD values were calculated for the morning and the evening windows. The larger the value, the greater the change in the activity waveform within that time window. The SSD values from all the individuals from a population were then averaged to yield the mean SSD for that population. Thus, resulting in 12 SSD values (*early_1-4_*, *control_1-4_* and *late_1-4_*) for morning as well as the evening window.

##### Change in the mean phase of activity (ΔφCoM)

The mean phase of activity was calculated within the morning and evening time windows from the normalised average activity profiles for the in-sync regime for an individual (Batschelet, 1981). The same calculations were carried out for the out-of-sync regime as well. Change in the mean phase of activity during the morning window was calculated by subtracting the value of the mean phase under out-of-sync regime from that of in-sync regime for a given individual. The same calculation was conducted for the evening window as well. The change in mean phase of activity within each time window from all the individuals from a population were then averaged to yield the average change in phase for that population. Thus, resulting in 12 Δφ_CoM_ values (*early_1-4_*, *control_1-4_* and *late_1-4_*) for morning as well as the evening window.

##### Statistical analysis

For both SSD and change in mean phase within each of the time windows for all six experiments, the twelve population values were used to conduct mixed model ANOVA with selection (*early*, *control* and *late*) and environmental regime as the fixed factors and blocks (1-4) as the random factor to assess differential responses of the two activity components (morning and evening) among the selection lines.

#### Time point sample collection for assaying *timeless* and *Psi* transcripts

3-5 days old virgin male flies from block 4 of *early*, *control,* and *late* selection lines were collected and housed in glass vials with corn-medium food. These vials were then placed in an incubator with a set environmental regime (MIR-154, Sanyo, Tokyo, Japan). The environmental regimes used in this study, along with the number of generations of selection before the common rearing for experimental flies, are tabulated below.

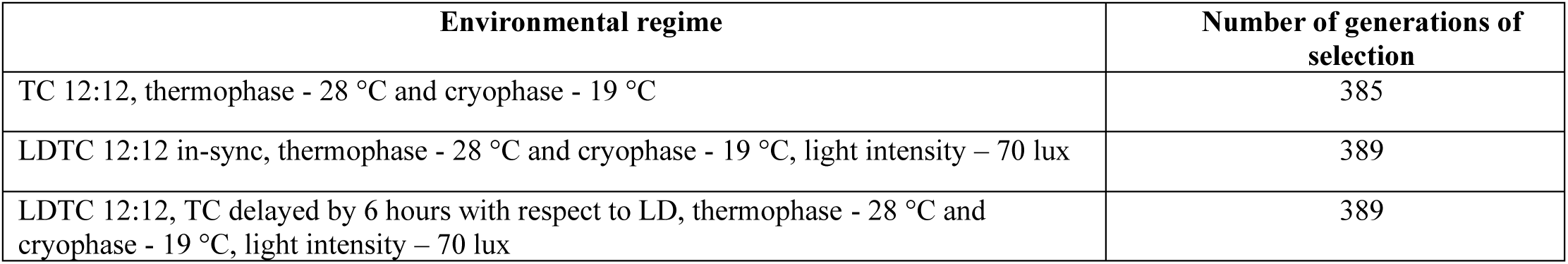

| Environmental regime | Number of generations of selection |
| --- | --- |
| TC 12:12, thermophase - 28 °C and cryophase - 19 °C | 385 |
| LDTC 12:12 in-sync, thermophase - 28 °C and cryophase - 19 °C, light intensity – 70 lux | 389 |
| LDTC 12:12, TC delayed by 6 hours with respect to LD, thermophase - 28 °C and cryophase - 19 °C, light intensity – 70 lux | 389 |

**Figure 11.**
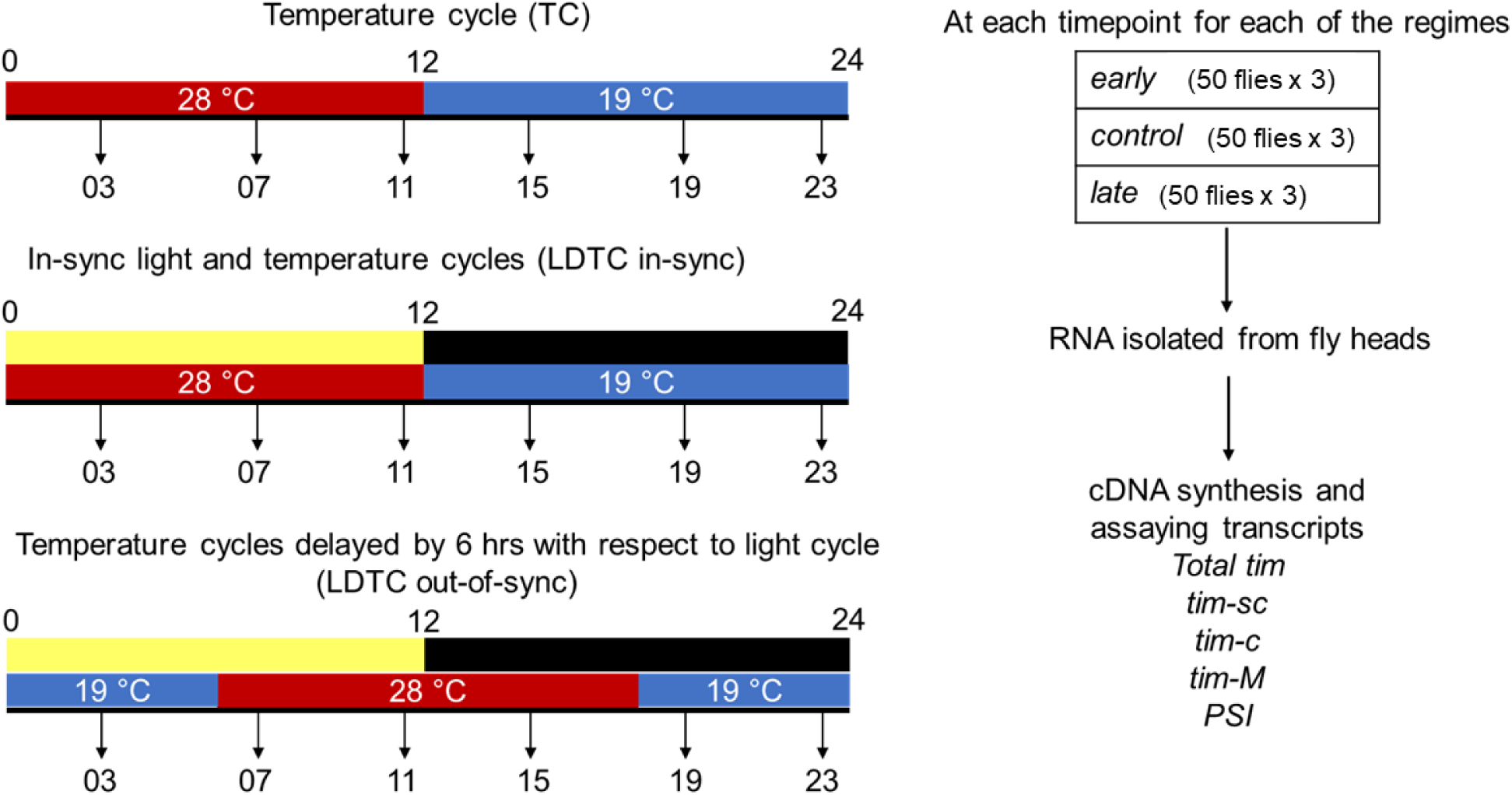
Summary of experimental protocol: *early_4_*, *control_4_* and *late_4_* flies (3-5 day old virgin males) after one generation of common rearing (no selection) were entrained to three different environmental regimes for 5 days. Sample collection at the above indicated time points were conducted on the 5^th^ cycle. Three biological replicates (with 50 flies each) were flash frozen using liquid nitrogen at each of the time point under all three regimes. Previously reported primers have been used in this study (see Table S6). Primers amplifying the exons 5&6 are used to quantify all *timeless* mRNA since these exons are common to all *timeless* variants. Amplification of the introns that are unique to each of the variants are used to quantify *tim-medium*, *tim-cold* and *tim-short and cold*.

### RNA extraction and quantitative real-time polymerase chain reaction

The flies were decapitated and at least 40-45 heads were used per sample to extract RNA. RNA was extracted using TRIzol/TRI reagent (Sigma-Alrich) as per the manufacturer’s instructions from the fly heads. 1 µg of this RNA was treated with DNase to remove any genomic DNA contamination. This served as the template for first-strand complementary DNA (cDNA) synthesis using the Maxima first-strand cDNA synthesis kit. Approximately 10-20 ng of cDNA was used in each qPCR reaction. The qPCR reactions were conducted in a Bio-rad CFX96 RT-PCR detection system. The qPCR mixture was subjected to 95 °C for 3 min, followed by 40 cycles of 95 °C for 10 s, 55 °C / 46 °C for 10 s, and 72 °C for 30 s, followed by a melting curve analysis to confirm that only one PCR product/peak is detected.

### Analyses of transcript expression profiles

The cycle threshold values obtained for each of the experimental transcripts were normalised by the expression of a housekeeping gene (*rp49/RpL32*) to obtain the relative expression of the experimental transcript for a biological replicate, for that time point, using the comparative Ct method (as per example 5 in Schmittgen & Livak, 2008). Each reaction was set up in duplicates and an average value of the two was used for the calculation of relative expression. Average expression and error across the three biological replicates have been plotted. The expression profiles have also been plotted with the expression level at each point scaled by the peak expression value and expressed as a percentage of the peak expression. This scaling was conducted for each biological replicate separately, thus facilitating both the visualisation of variation in expression levels across time points among biological replicates and comparison of profiles across regimes/experiments. These peak normalised expression profiles were used to quantify the mean phase of expression as well as the amplitude of expression (maximum – minimum expression level). RAIN analysis was used to detect rhythmicity in each dataset across a cycle (Thaben & Westermark, 2014). Two-way ANOVA tests with selection and time point as the factors were used to detect differences among the selection lines under a particular environmental regime. Amplitude and peak normalised expression patterns were used to test for the effect of environmental regime on the expression profile of a transcript in the wild-type *control* population followed by testing for differential effect of the environmental regime on the expression profiles among the selection lines (*early*, *control* and *late*).

GraphPad Prism v.8 was used for conducting all the one-way and two-way ANOVA tests. Mixed model ANOVA with three factors as well as repeated measures ANOVA were implemented in Statistica v.7.

Note: ‘expression pattern’, where used in this study refers only to the temporal expression pattern of the transcript across the time points under a given environmental regime.

All data was plotted using GraphPad Prism v.8

## Acknowledgements

We thank Abhilash Lakshman for experimental design in the early stages of the project and Shephali Dansana for carrying out initial pilot experiments which led to this project. We also thank research intern Prakruthi for carrying out RNA isolation for some of the samples. We would like to acknowledge financial support through intramural funding from the Jawaharlal Nehru Centre for Advanced Scientific Research (JNCASR). We also thank Rajanna, Muniraju and Samuel for their technical assistance.

## Author Contributions

Pragya Niraj Sharma., conceptualization, methodology, investigation, formal analysis, and writing-original draft and Sheeba Vasu., conceptualization, methodology, supervision, writing-review and editing, and funding acquisition.

## Supplementary information

**Figure S1.**
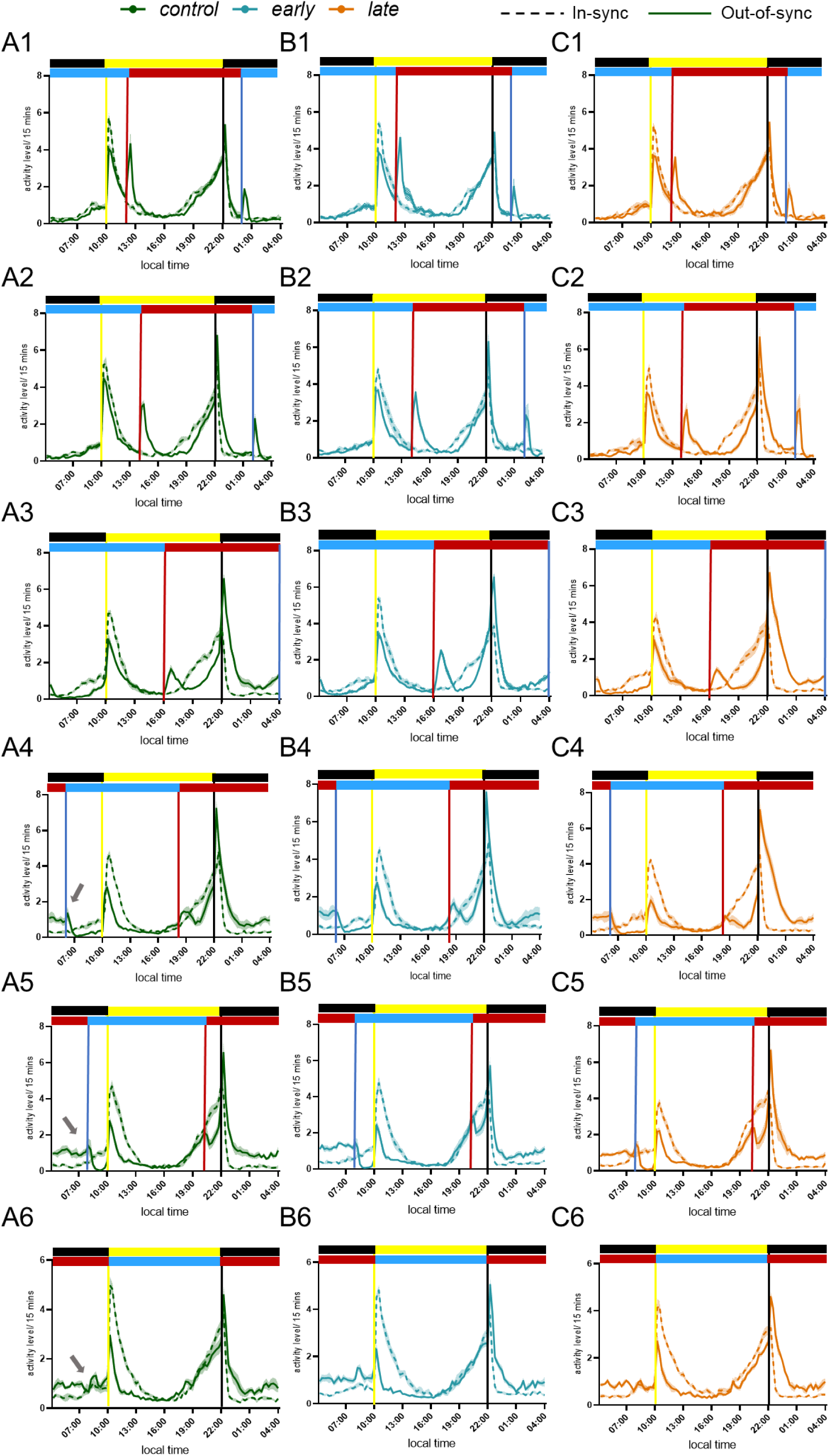
Activity profiles of divergent chronotype fly populations under environmental regimes with a range of phase relationship between the light and temperature cycles. (A1-6) Activity profiles of wildtype *Drosophila melanogaster* populations (*control_1-4_*) under environmental regimes with specific phase relationships between light and temperature cycles. (B1-6) Activity profiles of wildtype *Drosophila melanogaster* populations selected for morning fly emergence (*early_1-4_*) under environmental regimes with specific phase relationships between light and temperature cycles. (C1-6) Activity profiles of wildtype *Drosophila melanogaster* populations selected for evening fly emergence (*late_1-4_*) under environmental regimes with specific phase relationships between light and temperature cycles. Activity profiles under in-sync regime have been plotted as dashed traces while those under out-of-sync regimes have been plotted as solid traces for each of the experiments. Error bands are SEM across the four populations. The bars on top indicate the out-of-sync environmental regime with yellow and black indicating photophase and scotophase while red and blue indicate thermophase and cryophase, respectively.

**Figure S2.**
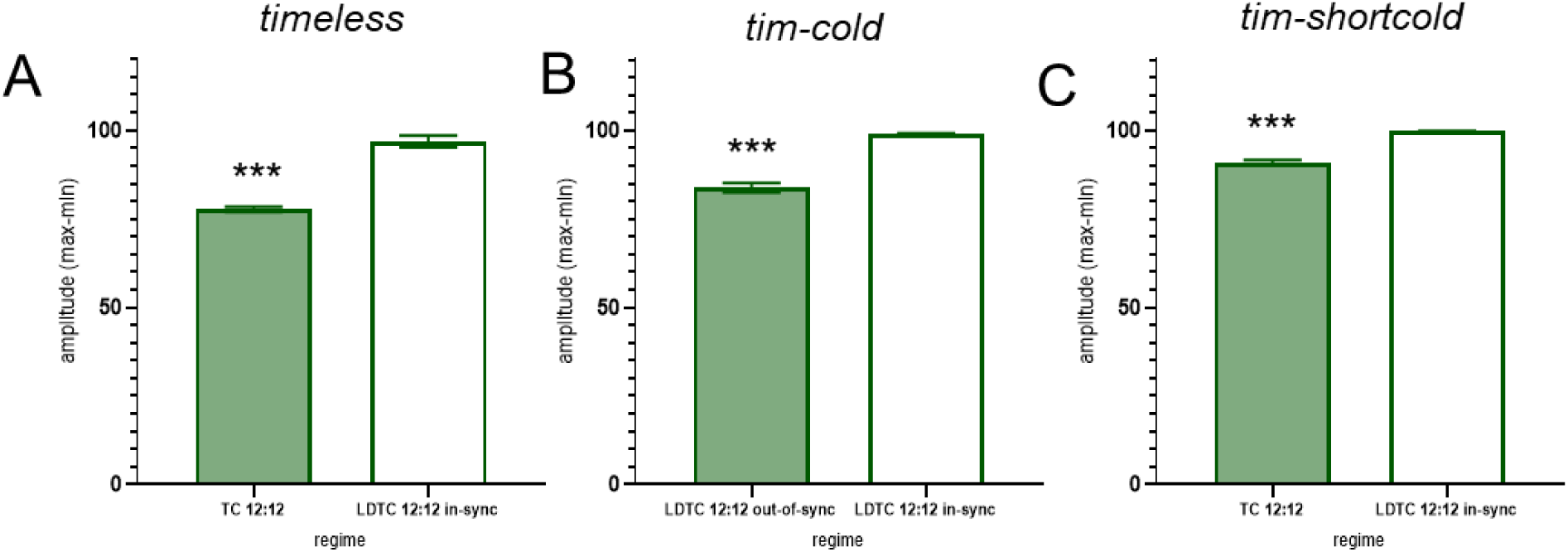
(A) Amplitude of *timeless* expression (maximum – minimum) compared between TC 12:12 and LDTC 12:12 in-sync regimes. The asterisks indicate that the amplitude was significantly lower under TC compared to LDTC in-sync conditions by an unpaired *t*-test (*p =* 0.0005). (B) Amplitude of *tim-cold* expression (maximum – minimum) compared between LDTC 12:12 out-of-sync and LDTC 12:12 in-sync regimes. The asterisks indicate that the amplitude was significantly lower under LDTC out-of-sync compared to LDTC in-sync conditions by an unpaired *t*-test (*p =* 0.0004). Error bars are SEM. (C) Amplitude of *tim-short and cold* expression (maximum – minimum) compared between TC 12:12 and LDTC 12:12 in-sync regimes. The asterisks indicate that the amplitude was significantly lower under TC compared to LDTC in-sync conditions by an unpaired *t*-test (*p =* 0.0003). Error bars are SEM.

**Figure S3.**
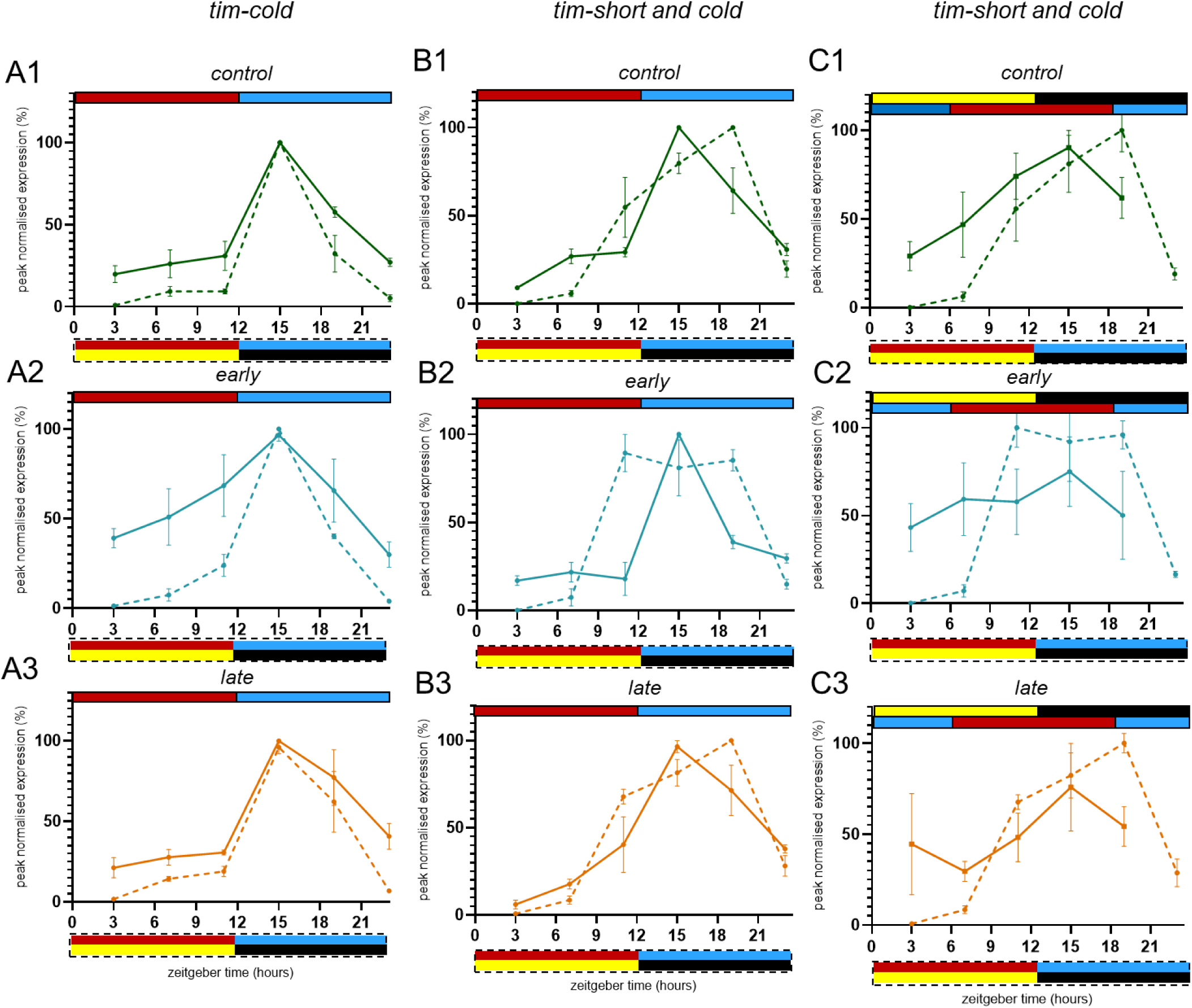
(A1-3) Peak normalised expression profile of *tim-cold* in control, early and late under TC 12:12 and under LDTC 12:12 in-sync conditions, solid and dashed traces, respectively. (B1-3) Peak normalised expression profile of *tim-sc* in *control*, *early* and *late* under TC 12:12 and under LDTC 12:12 in-sync conditions, solid and dashed traces, respectively. (C1-3) Peak normalised expression profile of *tim-sc* in control, early and late under LDTC out-of-sync and LDTC 12:12 in-sync conditions, solid and dashed traces, respectively. Environmental regimes for the solid and dashed traces are indicated at the top and bottom of the expression profiles, respectively. The bars indicating environmental regimes have solid or dashed outlines, corresponding to the solid or dashed traces for the respective expression profiles. Yellow and black bars for light and dark, respectively, while red and blue bars are for warm and cool phases, respectively. Error bars are SEM across three biological replicates.

**Figure S4.**
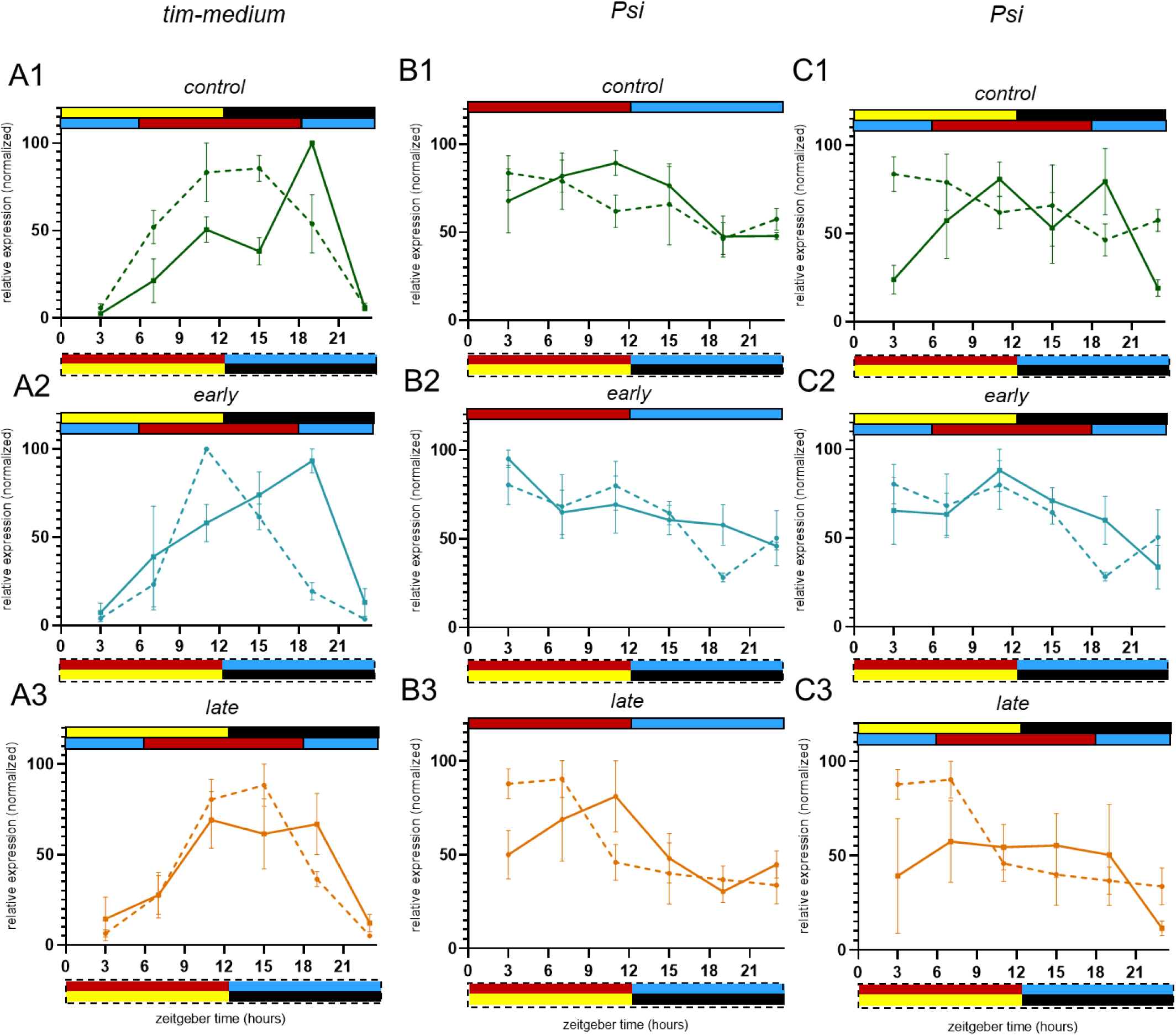
(A1-3) Peak normalised expression profile of *tim-medium* in *control*, *early* and *late* under TC 12:12 and under LDTC 12:12 in-sync conditions, solid and dashed traces, respectively. (B1-3) Peak normalised expression profile of *PSI* in *control*, *early* and *late* under TC 12:12 and under LDTC 12:12 in-sync conditions, solid and dashed traces, respectively. (C1-3) Peak normalised expression profile of *PSI* in *control*, *early* and *late* under LDTC out-of-sync and LDTC 12:12 in-sync conditions, solid and dashed traces, respectively. Environmental regimes for the solid and dashed traces are indicated at the top and bottom of the expression profiles, respectively. The bars indicating environmental regimes have solid or dashed outlines, corresponding to the solid or dashed traces for the respective expression profiles. Yellow and black bars for light and dark, respectively, while red and blue bars are for warm and cool phases, respectively. Error bars are SEM across three biological replicates.

**Table S1.** Two-way ANOVA tables for comparison of peak normalised expression values between two regimes for a given transcript. Regime and time point were the two factors, and the tables list their main effects as well as the interaction effect between the two factors. The transcript and the two environmental regimes between which its expression profile is compared are mentioned in the top cell of the first column for each table. For all the tables, statistically significant effects have been marked using red font.

|  |  |  |  |  |  |  |
| --- | --- | --- | --- | --- | --- | --- |
| A | **Total timeless TC & LDTC in-sync** | SS | DF | MS | F (DFn, DFd) | P value |
|  | Interaction | 8239 | 5 | 1648 | F (5, 24) = 7.139 | P=0.0003 |
|  | regime | 8199 | 1 | 8199 | F (1, 24) = 35.52 | P<0.0001 |
|  | time point | 22185 | 5 | 4437 | F (5, 24) = 19.22 | P<0.0001 |
|  | Residual | 5539 | 24 | 230.8 |  |  |
| B | **Total timeless LDTC in-sync & out-of-sync** | SS | DF | MS | F (DFn, DFd) | P value |
|  | Interaction | 5423 | 5 | 1085 | F (5, 24) = 2.986 | P=0.0310 |
|  | regime | 9508 | 1 | 9508 | F (1, 24) = 26.17 | P<0.0001 |
|  | time point | 18318 | 5 | 3664 | F (5, 24) = 10.08 | P<0.0001 |
|  | Residual | 8719 | 24 | 363.3 |  |  |
| C | **Tim-cold TC & LDTC in-sync** | SS | DF | MS | F (DFn, DFd) | P value |
|  | Interaction | 609.6 | 5 | 121.9 | F (5, 24) = 1.471 | P=0.2361 |
|  | regime | 2736 | 1 | 2736 | F (1, 24) = 33.01 | P<0.0001 |
|  | time point | 34799 | 5 | 6960 | F (5, 24) = 83.97 | P<0.0001 |
|  | Residual | 1989 | 24 | 82.89 |  |  |
| D | **Tim-cold LDTC in-sync & out-of-sync** | SS | DF | MS | F (DFn, DFd) | P value |
|  | Interaction | 15073 | 5 | 3015 | F (5, 24) = 50.42 | P<0.0001 |
|  | regime | 2562 | 1 | 2562 | F (1, 24) = 42.85 | P<0.0001 |
|  | time point | 19996 | 5 | 3999 | F (5, 24) = 66.89 | P<0.0001 |
|  | Residual | 1435 | 24 | 59.79 |  |  |
| E | **Tim-sc TC & LDTC in-sync** | SS | DF | MS | F (DFn, DFd) | P value |
|  | Interaction | 4473 | 5 | 894.5 | F (5, 24) = 6.456 | P=0.0006 |
|  | regime | 0.0371 | 1 | 0.03719 | F (1, 24) = 0.0002684 | P=0.9871 |
|  | time point | 37347 | 5 | 7469 | F (5, 24) = 53.91 | P<0.0001 |
|  | Residual | 3325 | 24 | 138.6 |  |  |
| F | **Tim-sc LDTC in-sync & out-of-sync** | SS | DF | MS | F (DFn, DFd) | P value |
|  | Interaction | 5537 | 4 | 1384 | F (4, 20) = 4.062 | P=0.0143 |
|  | regime | 1152 | 1 | 1152 | F (1, 20) = 3.380 | P=0.0809 |
|  | time point | 24708 | 4 | 6177 | F (4, 20) = 18.13 | P<0.0001 |
|  | Residual | 6815 | 20 | 340.8 |  |  |
| G | **Tim-medium TC & LDTC in-sync** | SS | DF | MS | F (DFn, DFd) | P value |
|  | Interaction | 3577 | 5 | 715.5 | F (5, 24) = 3.053 | P=0.0285 |
|  | regime | 8.392 | 1 | 8.392 | F (1, 24) = 0.03580 | P=0.8515 |
|  | time point | 33144 | 5 | 6629 | F (5, 24) = 28.28 | P<0.0001 |
|  | Residual | 5625 | 24 | 234.4 |  |  |
| H | **Tim-medium LDTC in-sync & out-of-sync** | SS | DF | MS | F (DFn, DFd) | P value |
|  | Interaction | 8410 | 5 | 1682 | F (5, 24) = 6.784 | P=0.0005 |
|  | regime | 1185 | 1 | 1185 | F (1, 24) = 4.778 | P=0.0388 |
|  | time point | 29888 | 5 | 5978 | F (5, 24) = 24.11 | P<0.0001 |
|  | Residual | 5950 | 24 | 247.9 |  |  |
| I | **Psi TC & LDTC in-sync** | SS | DF | MS | F (DFn, DFd) | P value |
|  | Interaction | 1756 | 5 | 351.1 | F (5, 24) = 0.7732 | P=0.5785 |
|  | regime | 69.04 | 1 | 69.04 | F (1, 24) = 0.1520 | P=0.7000 |
|  | time point | 5748 | 5 | 1150 | F (5, 24) = 2.531 | P=0.0563 |
|  | Residual | 10899 | 24 | 454.1 |  |  |
| J | **Psi LDTC in-sync & out-of-sync** | SS | DF | MS | F (DFn, DFd) | P value |
|  | Interaction | 9070 | 5 | 1814 | F (5, 24) = 2.912 | P=0.0341 |
|  | regime | 1637 | 1 | 1637 | F (1, 24) = 2.627 | P=0.1181 |
|  | time point | 4255 | 5 | 851.0 | F (5, 24) = 1.366 | P=0.2719 |
|  | Residual | 14951 | 24 | 622.9 |  |  |

**Table S2.** Three-way ANOVA tables with LDTC regime (phase relationship between light and temperature cycles) and selection as the main factors and blocks (4 populations that have undergone selection) as random factors. The measure (SSD or mean phase of activity) and the time window (morning or evening activity) is mentioned in the top cell of the first column.

|  |  |  |  |  |  |  |  |  |  |
| --- | --- | --- | --- | --- | --- | --- | --- | --- | --- |
| A | **SSD (morning window)** | Effect (F/R) | SS | Degr. of Freedom | MS | Den.Syn. Error df | Den.Syn. Error MS | F | p |
|  | Intercept | Fixed | 0.005673 | 1 | 0.005673 | 3.00000 | 0.000009 | 607.4123 | 0.000146 |
|  | regime | Fixed | 0.000218 | 5 | 0.000044 | 15.00000 | 0.000002 | 23.3916 | 0.000001 |
|  | selection | Fixed | 0.000038 | 2 | 0.000019 | 6.00000 | 0.000003 | 6.3821 | 0.032694 |
|  | block | Random | 0.000028 | 3 | 0.000009 | 7.66243 | 0.000004 | 2.5432 | 0.132604 |
|  | regime*selection | Fixed | 0.000043 | 10 | 0.000004 | 30.00000 | 0.000001 | 3.6466 | 0.002859 |
|  | regime*block | Random | 0.000028 | 15 | 0.000002 | 30.00000 | 0.000001 | 1.5874 | 0.137051 |
|  | selection*block | Random | 0.000018 | 6 | 0.000003 | 30.00000 | 0.000001 | 2.5345 | 0.041834 |
|  | regime*selection*block | Random | 0.000035 | 30 | 0.000001 | 0.00000 | 0.000000 |  |  |
|  | Error |  |  | 0 |  |  |  |  |  |
| B | **Mean phase ($\Delta\phi_{CoM}$) (morning window)** | Effect (F/R) | SS | Degr. of Freedom | MS | Den.Syn. Error df | Den.Syn. Error MS | F | p |
|  | Intercept | Fixed | 23.1774 | 1 | 23.17736 | 3.00000 | 0.547387 | 42.34182 | 0.007372 |
|  | regime | Fixed | 139.7711 | 5 | 27.95422 | 15.00000 | 0.282614 | 98.91321 | 0.000000 |
|  | selection | Fixed | 0.9621 | 2 | 0.48103 | 6.00000 | 0.275920 | 1.74338 | 0.252988 |
|  | block | Random | 1.6422 | 3 | 0.54739 | 10.73107 | 0.444884 | 1.23040 | 0.346094 |
|  | regime*selection | Fixed | 2.8852 | 10 | 0.28852 | 30.00000 | 0.113649 | 2.53867 | 0.023673 |
|  | regime*block | Random | 4.2392 | 15 | 0.28261 | 30.00000 | 0.113649 | 2.48673 | 0.016400 |
|  | selection*block | Random | 1.6555 | 6 | 0.27592 | 30.00000 | 0.113649 | 2.42783 | 0.049431 |
|  | regime*selection*block | Random | 3.4095 | 30 | 0.11365 | 0.00000 | 0.000000 |  |  |
|  | Error |  |  | 0 |  |  |  |  |  |
| C | **SSD (evening window)** | Effect (F/R) | SS | Degr. of Freedom | MS | Den.Syn. Error df | Den.Syn. Error MS | F | p |
|  | Intercept | Fixed | 0.020397 | 1 | 0.020397 | 3.00000 | 0.000386 | 52.77680 | 0.005382 |
|  | regime | Fixed | 0.007201 | 5 | 0.001440 | 15.00000 | 0.000227 | 6.33547 | 0.002368 |
|  | selection | Fixed | 0.000845 | 2 | 0.000423 | 6.00000 | 0.000054 | 7.83284 | 0.021239 |
|  | block | Random | 0.001159 | 3 | 0.000386 | 16.94668 | 0.000259 | 1.49420 | 0.251971 |
|  | regime*selection | Fixed | 0.000472 | 10 | 0.000047 | 30.00000 | 0.000023 | 2.08613 | 0.058543 |
|  | regime*block | Random | 0.003410 | 15 | 0.000227 | 30.00000 | 0.000023 | 10.04046 | 0.000000 |
|  | selection*block | Random | 0.000324 | 6 | 0.000054 | 30.00000 | 0.000023 | 2.38327 | 0.053008 |
|  | regime*selection*block | Random | 0.000679 | 30 | 0.000023 | 0.00000 | 0.000000 |  |  |
|  | Error |  |  | 0 |  |  |  |  |  |
| D | **Mean phase ($\Delta\phi_{CoM}$) (evening window)** | Effect (F/R) | SS | Degr. of Freedom | MS | Den.Syn. Error df | Den.Syn. Error MS | F | p |
|  | Intercept | Fixed | 70.01619 | 1 | 70.01619 | 3.00000 | 0.340406 | 205.6844 | 0.000735 |
|  | regime | Fixed | 6.53225 | 5 | 1.30645 | 15.00000 | 0.051244 | 25.4949 | 0.000001 |
|  | selection | Fixed | 7.99985 | 2 | 3.99992 | 6.00000 | 0.081167 | 49.2804 | 0.000189 |
|  | block | Random | 1.02122 | 3 | 0.34041 | 7.91716 | 0.101632 | 3.3494 | 0.076856 |
|  | regime*selection | Fixed | 1.88444 | 10 | 0.18844 | 30.00000 | 0.030778 | 6.1226 | 0.000051 |
|  | regime*block | Random | 0.76865 | 15 | 0.05124 | 30.00000 | 0.030778 | 1.6649 | 0.114401 |
|  | selection*block | Random | 0.48700 | 6 | 0.08117 | 30.00000 | 0.030778 | 2.6371 | 0.035652 |
|  | regime*selection*block | Random | 0.92335 | 30 | 0.03078 | 0.00000 | 0.000000 |  |  |
|  | Error |  |  | 0 |  |  |  |  |  |

**Table S3.**
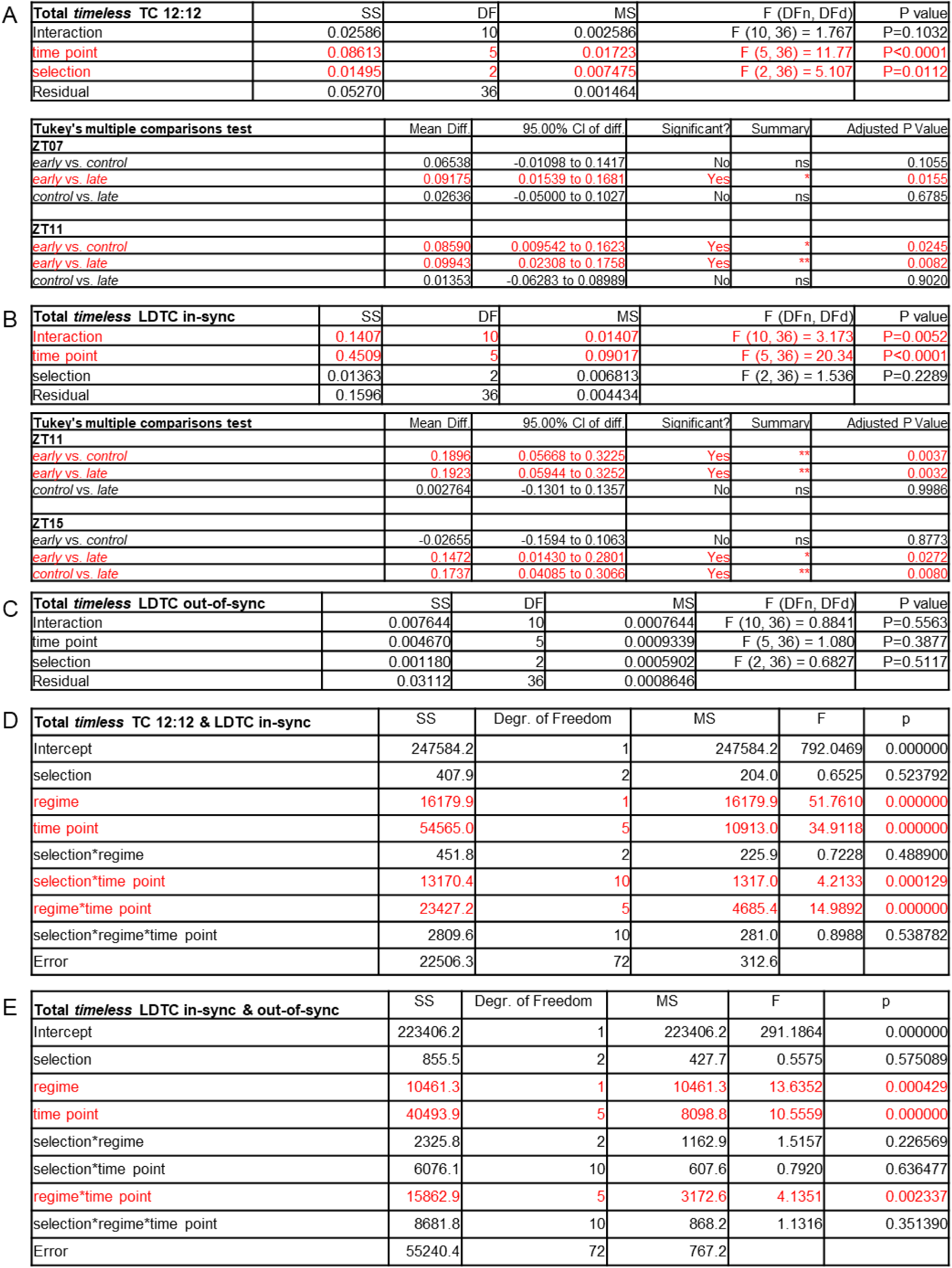
Expression profiles of total *timeless* in selection lines across three environmental regimes. (A) (top) Two-way ANOVA table comparing expression profiles of the transcript among the selection lines under TC 12:12. (bottom) Significant differences among the selection lines at indicated time points by Tukey’s multiple comparisons following the above two-way ANOVA. (B) (top) Two-way ANOVA table comparing expression profiles of the transcript among the selection lines under LDTC in-sync. (bottom) Significant differences among the selection lines at indicated time points by Tukey’s multiple comparisons following the above two-way ANOVA. (C) (top) Two-way ANOVA table comparing expression profiles of the transcript among the selection lines under LDTC out-of-sync. (D) Three-way ANOVA table for comparing expression profiles of the transcript across TC 12:12 and LDTC in-sync regimes among the selection lines. The table lists the three factors and their interaction effects. (D) Three-way ANOVA table for comparing expression profiles of the transcript across LDTC in-sync and LDTC out-of-sync regimes among the selection lines. The table lists the three factors and their interaction effects. For all the tables, statistically significant effects and differences have been marked by using red font.

| A | Total *timeless* TC 12:12 | SS | DF | MS | F (DFn, DFd) | P value |
| --- | --- | --- | --- | --- | --- | --- |
|  | Interaction | 0.02586 | 10 | 0.002586 | F (10, 36) = 1.767 | P=0.1032 |
|  | time point | 0.08613 | 5 | 0.01723 | F (5, 36) = 11.77 | P<0.0001 |
|  | selection | 0.01495 | 2 | 0.007475 | F (2, 36) = 5.107 | P=0.0112 |
|  | Residual | 0.05270 | 36 | 0.001464 |  |  |

| Tukey's multiple comparisons test | Mean Diff. | 95.00% CI of diff. | Significant? | Summary | Adjusted P Value |
| --- | --- | --- | --- | --- | --- |
| **ZT07** |  |  |  |  |  |
| early vs. control | 0.06538 | -0.01098 to 0.1417 | No | ns | 0.1055 |
| early vs. late | 0.09175 | 0.01539 to 0.1681 | Yes | * | 0.0155 |
| control vs. late | 0.02636 | -0.05000 to 0.1027 | No | ns | 0.6785 |
| **ZT11** |  |  |  |  |  |
| early vs. control | 0.08590 | 0.009542 to 0.1623 | Yes | * | 0.0245 |
| early vs. late | 0.09943 | 0.02308 to 0.1758 | Yes | ** | 0.0082 |
| control vs. late | 0.01353 | -0.06283 to 0.08989 | No | ns | 0.9020 |

| B | Total *timeless* LDTC in-sync | SS | DF | MS | F (DFn, DFd) | P value |
| --- | --- | --- | --- | --- | --- | --- |
|  | Interaction | 0.1407 | 10 | 0.01407 | F (10, 36) = 3.173 | P=0.0052 |
|  | time point | 0.4509 | 5 | 0.09017 | F (5, 36) = 20.34 | P<0.0001 |
|  | selection | 0.01363 | 2 | 0.006813 | F (2, 36) = 1.536 | P=0.2289 |
|  | Residual | 0.1596 | 36 | 0.004434 |  |  |

| Tukey's multiple comparisons test | Mean Diff. | 95.00% CI of diff. | Significant? | Summary | Adjusted P Value |
| --- | --- | --- | --- | --- | --- |
| **ZT11** |  |  |  |  |  |
| early vs. control | 0.1896 | 0.05668 to 0.3225 | Yes | ** | 0.0037 |
| early vs. late | 0.1923 | 0.05944 to 0.3252 | Yes | ** | 0.0032 |
| control vs. late | 0.002764 | -0.1301 to 0.1357 | No | ns | 0.9986 |
| **ZT15** |  |  |  |  |  |
| early vs. control | -0.02655 | -0.1594 to 0.1063 | No | ns | 0.8773 |
| early vs. late | 0.1472 | 0.01430 to 0.2801 | Yes | * | 0.0272 |
| control vs. late | 0.1737 | 0.04085 to 0.3066 | Yes | ** | 0.0080 |

| C | Total *timeless* LDTC out-of-sync | SS | DF | MS | F (DFn, DFd) | P value |
| --- | --- | --- | --- | --- | --- | --- |
|  | Interaction | 0.007644 | 10 | 0.0007644 | F (10, 36) = 0.8841 | P=0.5563 |
|  | time point | 0.004670 | 5 | 0.0009339 | F (5, 36) = 1.080 | P=0.3877 |
|  | selection | 0.001180 | 2 | 0.0005902 | F (2, 36) = 0.6827 | P=0.5117 |
|  | Residual | 0.03112 | 36 | 0.0008646 |  |  |

| D | Total *timeless* TC 12:12 & LDTC in-sync | SS | Degr. of Freedom | MS | F | p |
| --- | --- | --- | --- | --- | --- | --- |
|  | Intercept | 247584.2 | 1 | 247584.2 | 792.0469 | 0.000000 |
|  | selection | 407.9 | 2 | 204.0 | 0.6525 | 0.523792 |
|  | regime | 16179.9 | 1 | 16179.9 | 51.7610 | 0.000000 |
|  | time point | 54565.0 | 5 | 10913.0 | 34.9118 | 0.000000 |
|  | selection*regime | 451.8 | 2 | 225.9 | 0.7228 | 0.488900 |
|  | selection*time point | 13170.4 | 10 | 1317.0 | 4.2133 | 0.000129 |
|  | regime*time point | 23427.2 | 5 | 4685.4 | 14.9892 | 0.000000 |
|  | selection*regime*time point | 2809.6 | 10 | 281.0 | 0.8988 | 0.538782 |
|  | Error | 22506.3 | 72 | 312.6 |  |  |

| E | Total *timeless* LDTC in-sync & out-of-sync | SS | Degr. of Freedom | MS | F | p |
| --- | --- | --- | --- | --- | --- | --- |
|  | Intercept | 223406.2 | 1 | 223406.2 | 291.1864 | 0.000000 |
|  | selection | 855.5 | 2 | 427.7 | 0.5575 | 0.575089 |
|  | regime | 10461.3 | 1 | 10461.3 | 13.6352 | 0.000429 |
|  | time point | 40493.9 | 5 | 8098.8 | 10.5559 | 0.000000 |
|  | selection*regime | 2325.8 | 2 | 1162.9 | 1.5157 | 0.226569 |
|  | selection*time point | 6076.1 | 10 | 607.6 | 0.7920 | 0.636477 |
|  | regime*time point | 15862.9 | 5 | 3172.6 | 4.1351 | 0.002337 |
|  | selection*regime*time point | 8681.8 | 10 | 868.2 | 1.1316 | 0.351390 |
|  | Error | 55240.4 | 72 | 767.2 |  |  |

**Table S4.** Expression profiles of cold-induced *timeless* transcripts in selection lines across three environmental regimes. (A) Two-way ANOVA table comparing expression profiles of *tim-cold* among the selection lines under TC 12:12. (B) Two-way ANOVA table comparing expression profiles of *tim-cold* among the selection lines under LDTC in-sync. (C) Two-way ANOVA table comparing expression profiles of *tim-cold* among the selection lines under LDTC out-of-sync. (D) Three-way ANOVA table for comparing expression profiles of *tim-cold* across TC 12:12 and LDTC in-sync regimes among the selection lines. The table lists the three factors and their interaction effects. (E) Three-way ANOVA table for comparing expression profiles of *tim-cold* across LDTC in-sync and LDTC out-of-sync regimes among the selection lines. The table lists the three factors and their interaction effects. (F) Two-way ANOVA table comparing expression profiles of *tim-short and cold* among the selection lines under TC 12:12. (G) Two-way ANOVA table comparing expression profiles of *tim-short and cold* among the selection lines under LDTC in-sync. (H) Two-way ANOVA table comparing expression profiles of *tim-short and cold* among the selection lines under LDTC out-of-sync. (I) Three-way ANOVA table for comparing expression profiles of *tim-short and cold* across TC 12:12 and LDTC in-sync regimes among the selection lines. The table lists the three factors and their interaction effects. (J) Three-way ANOVA table for comparing expression profiles of *tim-short and cold* across LDTC in-sync and LDTC out-of-sync regimes among the selection lines. The table lists the three factors and their interaction effects. For all the tables, statistically significant effects and differences have been marked by using red font.

|  |  |  |  |  |  |  |
| --- | --- | --- | --- | --- | --- | --- |
| A | **Tim-cold TC 12:12** | SS | DF | MS | F (DFn, DFd) | P value |
|  | Interaction | 0.003784 | 10 | 0.0003784 | F (10, 36) = 1.936 | P=0.0720 |
|  | time point | 0.02870 | 5 | 0.005740 | F (5, 36) = 29.37 | P<0.0001 |
|  | selection | 0.0009128 | 2 | 0.0004564 | F (2, 36) = 2.335 | P=0.1113 |
|  | Residual | 0.007036 | 36 | 0.0001954 |  |  |
| B | **Tim-cold LDTC in-sync** | SS | DF | MS | F (DFn, DFd) | P value |
|  | Interaction | 0.02156 | 10 | 0.002156 | F (10, 36) = 1.249 | P=0.2949 |
|  | time point | 0.1678 | 5 | 0.03355 | F (5, 36) = 19.44 | P<0.0001 |
|  | selection | 0.003843 | 2 | 0.001921 | F (2, 36) = 1.113 | P=0.3395 |
|  | Residual | 0.06214 | 36 | 0.001726 |  |  |
| C | **Tim-cold LDTC out-of-sync** | SS | DF | MS | F (DFn, DFd) | P value |
|  | Interaction | 0.0003347 | 10 | 3.347e-005 | F (10, 36) = 0.9964 | P=0.4646 |
|  | time point | 0.004954 | 5 | 0.0009908 | F (5, 36) = 29.49 | P<0.0001 |
|  | selection | 7.124e-005 | 2 | 3.562e-005 | F (2, 36) = 1.060 | P=0.3569 |
|  | Residual | 0.001209 | 36 | 3.359e-005 |  |  |
| D | **Tim-cold TC 12:12 & LDTC in-sync** | SS | Degr. of Freedom | MS | F | p |
|  | Intercept | 174010.4 | 1 | 174010.4 | 973.7861 | 0.000000 |
|  | selection | 1574.8 | 2 | 787.4 | 4.4065 | 0.015655 |
|  | regime | 11773.9 | 1 | 11773.9 | 65.8883 | 0.000000 |
|  | time point | 93837.6 | 5 | 18767.5 | 105.0255 | 0.000000 |
|  | selection*regime | 882.7 | 2 | 441.4 | 2.4699 | 0.091733 |
|  | selection*time point | 3646.9 | 10 | 364.7 | 2.0409 | 0.041130 |
|  | regime*time point | 2379.3 | 5 | 475.9 | 2.6630 | 0.028930 |
|  | selection*regime*time point | 1379.2 | 10 | 137.9 | 0.7718 | 0.655147 |
|  | Error | 12866.0 | 72 | 178.7 |  |  |
| E | **Tim-cold LDTC in-sync & LDTC out-of-sync** | SS | Degr. of Freedom | MS | F | p |
|  | Intercept | 153918.1 | 1 | 153918.1 | 1325.211 | 0.000000 |
|  | selection | 562.0 | 2 | 281.0 | 2.419 | 0.096181 |
|  | regime | 7003.3 | 1 | 7003.3 | 60.298 | 0.000000 |
|  | time point | 63936.0 | 5 | 12787.2 | 110.096 | 0.000000 |
|  | selection*regime | 107.6 | 2 | 53.8 | 0.463 | 0.631206 |
|  | selection*time point | 1632.9 | 10 | 163.3 | 1.406 | 0.195244 |
|  | regime*time point | 38305.9 | 5 | 7661.2 | 65.962 | 0.000000 |
|  | selection*regime*time point | 2205.7 | 10 | 220.6 | 1.899 | 0.059140 |
|  | Error | 8362.5 | 72 | 116.1 |  |  |
| F | **Tim-sc TC 12:12** | SS | DF | MS | F (DFn, DFd) | P value |
|  | Interaction | 0.0005643 | 10 | 5.643e-005 | F (10, 36) = 1.319 | P=0.2576 |
|  | time point | 0.01028 | 5 | 0.002056 | F (5, 36) = 48.05 | P<0.0001 |
|  | selection | 0.0001197 | 2 | 5.987e-005 | F (2, 36) = 1.400 | P=0.2598 |
|  | Residual | 0.001540 | 36 | 4.278e-005 |  |  |
| G | **Tim-sc LDTC in-sync** | SS (Type III) | DF | MS | F (DFn, DFd) | P value |
|  | Interaction | 0.001139 | 10 | 0.0001139 | F (10, 35) = 1.273 | P=0.2829 |
|  | time point | 0.02294 | 5 | 0.004588 | F (5, 35) = 51.27 | P<0.0001 |
|  | selection | 0.0003212 | 2 | 0.0001606 | F (2, 35) = 1.795 | P=0.1811 |
|  | Residual | 0.003132 | 35 | 8.949e-005 |  |  |
| H | **Tim-sc LDTC out-of-sync** | SS (Type III) | DF | MS | F (DFn, DFd) | P value |
|  | Interaction | 6.984e-005 | 8 | 8.730e-006 | F (8, 29) = 0.1535 | P=0.9953 |
|  | time point | 0.0006765 | 4 | 0.0001691 | F (4, 29) = 2.974 | P=0.0357 |
|  | selection | 5.163e-005 | 2 | 2.581e-005 | F (2, 29) = 0.4538 | P=0.6396 |
|  | Residual | 0.001649 | 29 | 5.688e-005 |  |  |
| I | **Tim-sc TC 12:12 & LDTC in-sync** | SS | Degr. of Freedom | MS | F | p |
|  | Intercept | 216843.2 | 1 | 216843.2 | 963.3654 | 0.000000 |
|  | selection | 165.6 | 2 | 82.8 | 0.3679 | 0.693458 |
|  | regime | 844.2 | 1 | 844.2 | 3.7506 | 0.056714 |
|  | time point | 99872.2 | 5 | 19974.4 | 88.7401 | 0.000000 |
|  | selection*regime | 1004.1 | 2 | 502.0 | 2.2304 | 0.114866 |
|  | selection*time point | 3801.9 | 10 | 380.2 | 1.6891 | 0.099903 |
|  | regime*time point | 15675.5 | 5 | 3135.1 | 13.9282 | 0.000000 |
|  | selection*regime*time point | 2528.2 | 10 | 252.8 | 1.1232 | 0.357305 |
|  | Error | 16206.4 | 72 | 225.1 |  |  |
| J | **Tim-sc LDTC in-sync & out-of-sync** | SS | Degr. of Freedom | MS | F | p |
|  | Intercept | 275002.1 | 1 | 275002.1 | 403.4282 | 0.000000 |
|  | selection | 348.3 | 2 | 174.2 | 0.2555 | 0.775368 |
|  | regime | 496.8 | 1 | 496.8 | 0.7288 | 0.396667 |
|  | time point | 50497.4 | 4 | 12624.4 | 18.5199 | 0.000000 |
|  | selection*regime | 774.1 | 2 | 387.1 | 0.5678 | 0.569774 |
|  | selection*time point | 3076.1 | 8 | 384.5 | 0.5641 | 0.802968 |
|  | regime*time point | 19468.1 | 4 | 4867.0 | 7.1399 | 0.000090 |
|  | selection*regime*time point | 2439.3 | 8 | 304.9 | 0.4473 | 0.887529 |
|  | Error | 40899.8 | 60 | 681.7 |  |  |

**Table S5.**
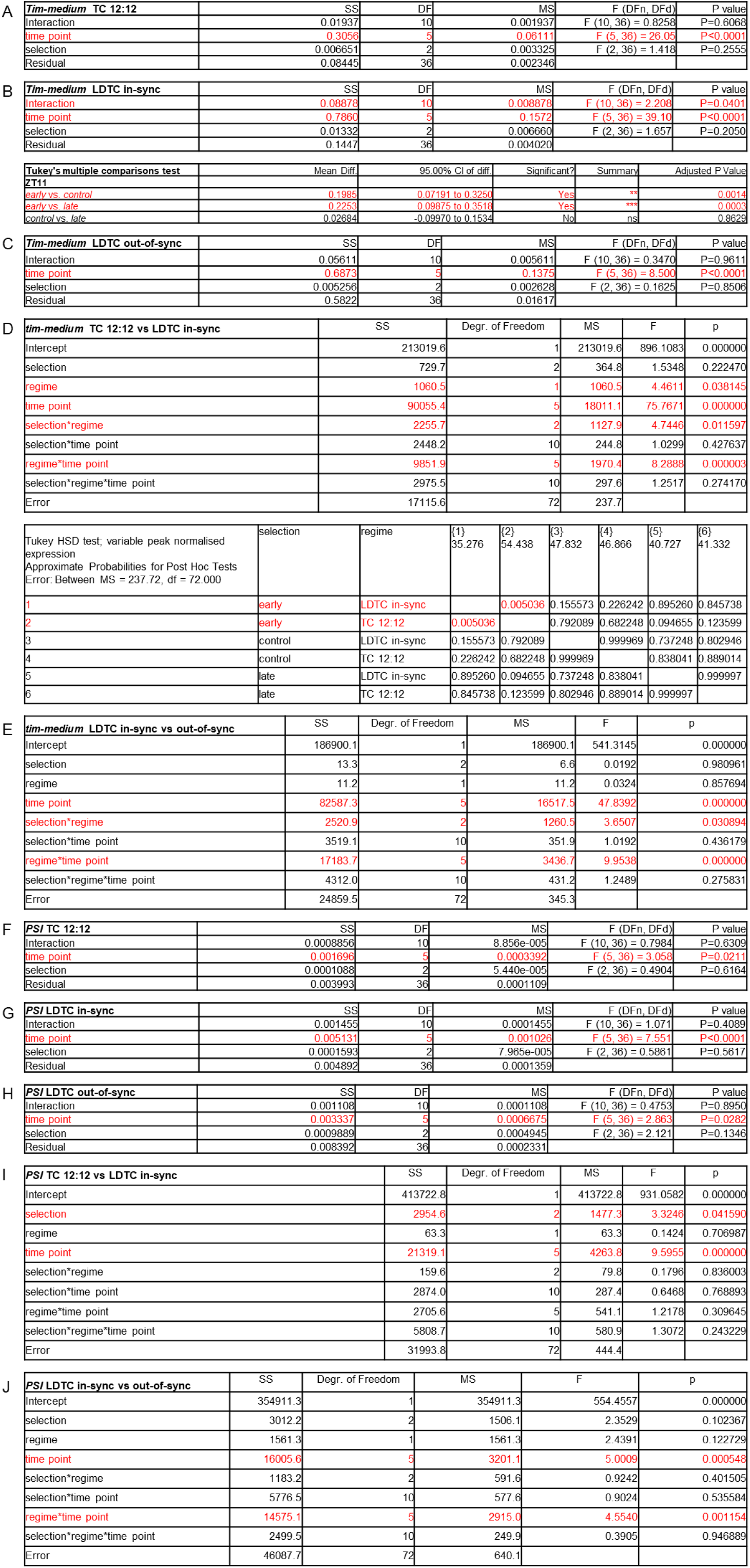
Expression profiles of warm-induced *timeless* transcript *tim-medium* and *timeless* splicing regulator *Psi* in selection lines across three environmental regimes. (A) Two-way ANOVA table comparing expression profiles of *tim-medium* among the selection lines under TC 12:12. (B) Two-way ANOVA table comparing expression profiles of *tim-medium* among the selection lines under LDTC in-sync. (bottom) Significant differences among the selection lines at the indicated time point by Tukey’s multiple comparisons following the above two-way ANOVA. (C) Two-way ANOVA table comparing expression profiles of *tim-medium* among the selection lines under LDTC out-of-sync. (D) Three-way ANOVA table for comparing expression profiles of *tim-medium* across TC 12:12 and LDTC in-sync regimes among the selection lines. The table lists the three factors and their interaction effects. (bottom) Significant differences from Tukey’s HSD following the above ANOVA for the interaction term between selection and regime. (E) Three-way ANOVA table for comparing expression profiles of *tim-medium* across LDTC in-sync and LDTC out-of-sync regimes among the selection lines. The table lists the three factors and their interaction effects. (F) Two-way ANOVA table comparing expression profiles of *Psi* among the selection lines under TC 12:12. (G) Two-way ANOVA table comparing expression profiles of *Psi* among the selection lines under LDTC in-sync. (H) Two-way ANOVA table comparing expression profiles of *Psi* among the selection lines under LDTC out-of-sync. (I) Three-way ANOVA table for comparing expression profiles of *Psi* across TC 12:12 and LDTC in-sync regimes among the selection lines. The table lists the three factors and their interaction effects. (J) Three-way ANOVA table for comparing expression profiles of *Psi* across LDTC in-sync and LDTC out-of-sync regimes among the selection lines. The table lists the three factors and their interaction effects. For all the tables, statistically significant effects and differences have been marked using red font.

**Table S6.**
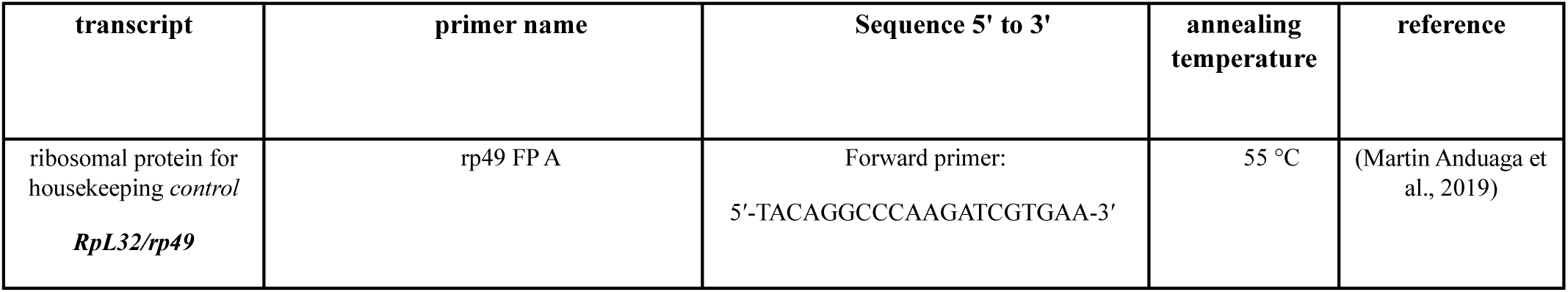
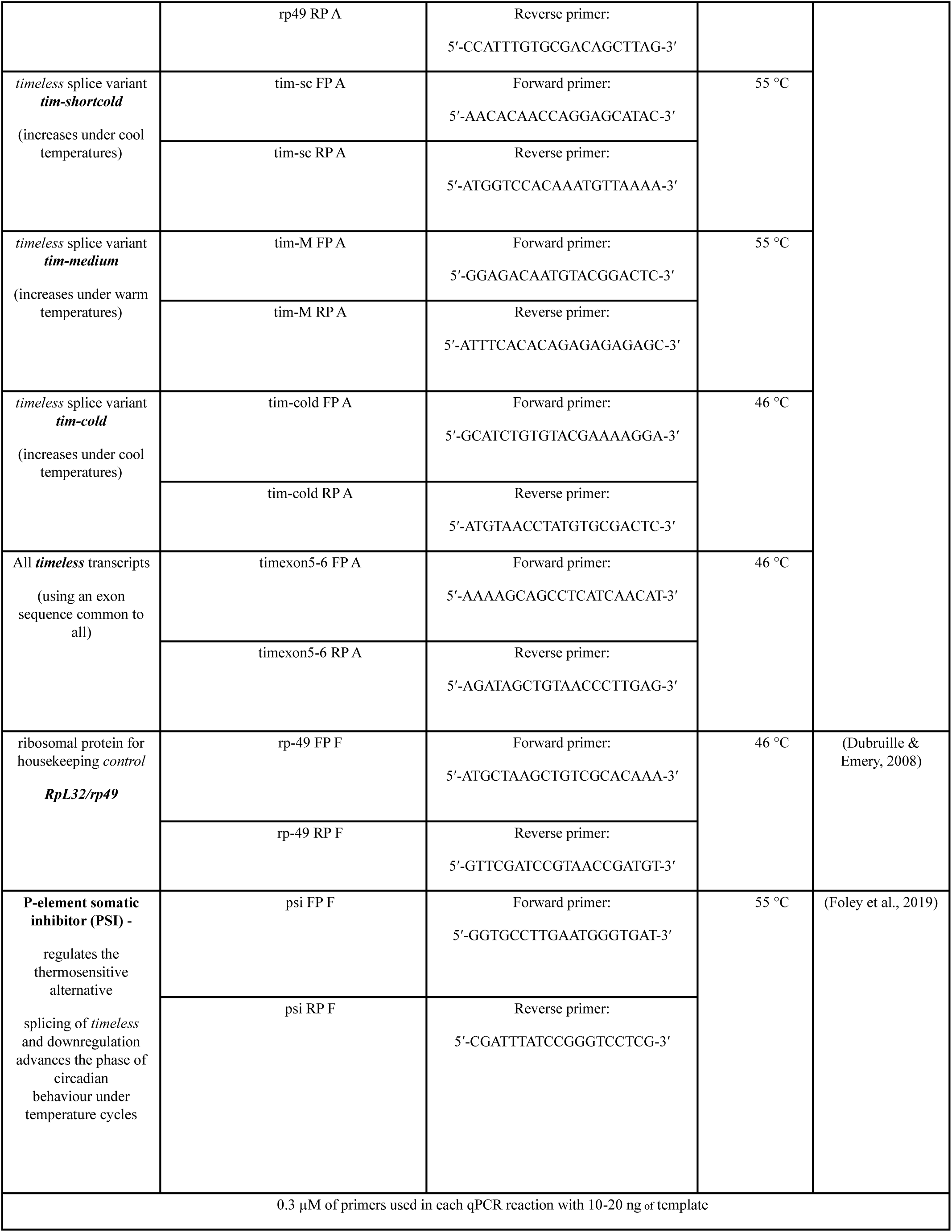
Details of all the primers used in this study along with their sequences, annealing temperatures used in this study and references.

